# Stress-induced Membrane Insertion at the β-Barrel Assembly Machinery Complex Regulates BepA Metalloprotease Activity

**DOI:** 10.64898/2026.07.01.735768

**Authors:** Henri Voedts, Phuong Chi Nguyen, Van-Son Nguyen, Pauline Leverrier, Bogdan Iorga, Seung Hyun Cho, Han Remaut, Jean-François Collet

**Affiliations:** WELBIO department, WEL Research Institute, avenue Pasteur, 6, 1300 Wavre, Belgium; de Duve Institute, Université catholique de Louvain (UCLouvain), Avenue Hippocrate 75, 1200 Brussels, Belgium; Structural Biology Brussels, Vrije Universiteit Brussel, 1050 Brussels, Belgium; Structural and Molecular Microbiology, Structural Biology Research Center, VIB, 1050 Brussels, Belgium; Université Paris-Saclay, CNRS UPR 2301, Institut de Chimie des Substances Naturelles, 91198 Gif-sur-Yvette, France

## Abstract

Proteases must be tightly regulated to prevent uncontrolled degradation, yet the mechanisms ensuring such control remain poorly understood. Members of the widespread M48 metalloprotease family are kept inactive by an autoinhibitory plug that blocks catalytic water activation, but how this plug is released was unknown. Here, using genetic, biochemical and cryo-EM approaches, we discover the activation mechanism of BepA, a quality-control protease that preserves outer membrane integrity by surveilling the β-barrel assembly machinery (BAM) in Gram-negative bacteria. Our cryo-EM analysis of BepA engaged with a stalled BAM-substrate assembly complex revealed that a flexible, unstructured α6-lid covering the active site in the latent protein functions as a molecular harpoon, inserting into the outer membrane when a substrate stalls at BAM and thereby docking BepA at the complex. This membrane anchoring promotes displacement of the autoinhibitory plug and unlocks protease activity precisely where and when it is needed. Thus, a dual enzyme activation mechanism is coupled to membrane association under stress, ensuring that BepA remains inactive until properly localized. Our findings reveal how membranes themselves can license protease activation, a principle that may extend beyond M48 metalloproteases.

## INTRODUCTION

The cell envelope is the morphological hallmark of Gram-negative bacteria. It consists of two membranes that enclose the periplasm, an aqueous, oxidizing compartment containing the peptidoglycan cell wall^1^. The outer membrane plays a crucial role in maintaining cell integrity by acting both as a permeability barrier against toxic compounds^2^ and as a load-bearing structure that supports periplasmic turgor buildup^3,4^. Two major classes of proteins reside within the outer membrane: lipoproteins, which are globular proteins tethered to the membrane by a lipid moiety, and outer membrane proteins (OMPs), which are integral membrane proteins with a β-barrel conformation^1^. OMPs are synthesized in the cytoplasm and are secreted, unfolded, into the periplasm via the Sec pathway. They are then escorted to the outer membrane by periplasmic chaperones, which deliver them to the β-barrel Assembly Machinery (BAM) for insertion into the outer membrane^5^. BamA, the central component of BAM, is a 16-stranded β-barrel protein belonging to the conserved OMP85 superfamily^6^. At its N-terminus, BamA displays five polypeptide transport-associated (POTRA) domains extending into the periplasm. In *Escherichia coli*, the BAM holocomplex comprises BamA and four accessory lipoproteins (BamB, BamC, BamD, and BamE), which interact with BamA’s POTRA domains. Although only BamA and BamD are essential for viability, all BAM subunits are required for optimal complex function.

Recent structural studies have begun to elucidate how BAM mediates OMP’s assembly^7–11^. Central to this process is the conformational flexibility of BamA, which alternates between two distinct states: an inward-open conformation, where the lateral seam formed by β-strands β1 and β16 remains sealed, and an outward-open conformation, in which this seam opens to template substrate folding and membrane insertion. Cycling between these two conformations is essential for the insertion and folding of OMP substrates. Current evidence indicates that OMP folding is initiated through binding of a conserved sequence at the C-terminal β-strand of a substrate (the so-called “β-signal”) to the β1 strand of BamA. Subsequent sequential addition of β-strands from the substrate’s C-terminus toward its N-terminus results in the formation of an asymmetric hybrid barrel, where BamA adopts its outward-open conformation. Facilitated by outer membrane tension and membrane destabilization induced by BamA, the substrate β-barrel is spawning off the exposed β1 seam of BamA and ultimately released into the outer membrane. During the assembly cycle, the POTRA domains undergo structural rearrangements. Positioned underneath the barrel and extending into the periplasm, they block access to the BamA β-barrel’s lumen when the lateral gate is open, but shift away when the seam closes, thereby creating a periplasmic entry pore into the barrel lumen.

OMP processing by BAM is not foolproof. Misfolding of OMPs can result from environmental stresses that disrupt BAM function or from intrinsic defects in the assembly pathway, such as stalling at the level of BamA. These stalled substrates risk clogging the BAM complex, leading to accumulation of misfolded OMPs in the periplasm and compromised envelope integrity^5^. To counteract these threats, bacteria have evolved quality control systems that continuously monitor OMP assembly and mount a protective response when envelope integrity is at risk ^12–14^. One key system is the σ^E^ stress response, which detects the accumulation of misfolded OMPs in the periplasm and triggers the upregulation of chaperones and proteases to restore envelope proteostasis^15–17^. The σ^E^ regulon includes BepA, a periplasmic, soluble quality control factor with an N-terminal protease domain from the M48 metallopeptidase family and a C-terminal tetratricopeptide repeat (TPR) domain (Extended Data Figure 1a)^18–21^. M48 metalloproteases are zinc-dependent enzymes widely distributed across bacteria and eukaryotes, where they play key roles in membrane-associated protein turnover^22–28^. BepA interacts with the BAM complex via its TPR domain and plays an important role in the assembly of LptD, an essential β-barrel responsible for LPS insertion into the outer membrane^18,19^. It has a dual function: under normal conditions, BepA promotes LptD assembly, but when folding fails, it switches to degrade misassembled LptD trapped at BAM^18,29,30^. Accordingly, in the absence of BepA, immature LptD accumulates in the outer membrane, leading to increased permeability to large antibiotics such as vancomycin^18,21^. BepA may also target additional stalled substrates at BAM, as suggested by its ability to degrade misfolded BamA under envelope stress^18,21^.

BepA displays very low proteolytic activity *in vitro*^18^, and the protease is thought to become activated specifically in the context of OMP assembly failures, targeting substrates that are engaged with the BAM complex but stalled during folding^5^. Interestingly, crystal structures of BepA revealed that a segment within its protease domain (residues R^234^ to P^249^, containing α9, Extended Data Figures 1a and 1b) acts as an active-site autoinhibitory-plug (AI-plug), occluding the catalytic residues and maintaining the protein in an autoinhibited state^21,29^. In this conformation, a conserved histidine residue (H^246^) on the AI-plug coordinates the active-site Zn^2+^ ion, effectively blocking the catalytic water molecule required for peptide-bond hydrolysis. This histidine serves as a reversible “off-switch”: BepA variants lacking H^246^ exhibit increased proteolytic activity. However, the molecular signals that trigger disengagement of the AI-plug and displacement of H^246^ to activate BepA remain unclear. Additionally, the mechanisms that regulate BepA’s dynamic association with the outer membrane-embedded BAM complex under stress conditions to clear stalled substrates are not yet fully understood.

Given the critical role of BepA in safeguarding outer membrane integrity and the unresolved questions surrounding how its protease activity is precisely regulated to function only when and where needed, we set out to elucidate the molecular mechanism underlying BepA activation and its coordination with OMP biogenesis in BAM. Using a combination of bacterial genetics and biochemical approaches, we found that the movement of a flexible α6-lid covering the active site is a prerequisite for the displacement of the AI-plug, a key step in BepA activation. Cryo-EM analysis of a BepA-BAM-substrate complex revealed that α6-lid movement involves its insertion into the outer membrane, near the substrate folding site at the lateral seam of BamA; thus, the flexible α6-lid functions as a unique molecular harpoon, triggered when OMP folding stalls, anchoring BepA to the membrane and stabilizing its interaction with BAM. This spatially controlled mechanism ensures that BepA remains inactive in the periplasm, thereby protecting the cell from unwanted proteolysis, and becomes proteolytically competent only upon recruitment to the outer membrane and engagement with stalled OMP substrates at BAM.

## RESULTS

### Identification of BepA variants impairing outer membrane integrity

To gain molecular insights into the mechanism of BepA and its regulation, we performed an unbiased genetic screen to identify key residues critical for its function. Given that BepA dysfunction compromises outer membrane integrity^18^, we generated a plasmid library of *bepA* alleles through random PCR mutagenesis and introduced them into Δ*bepA* cells. Transformants were then screened for increased outer membrane permeability using chlorophenyl red-β-D-galactopyranoside (CPRG). This assay exploits the impermeability of the intact outer membrane to CPRG, a β-galactosidase substrate that produces a pink color upon hydrolysis. Transformants with increased outer membrane permeability allow CPRG to enter the cell, where it is hydrolyzed, producing a detectable pink color^31^ (Figure 1a). Pink transformants were isolated, their plasmids extracted and reintroduced into Δ*bepA* cells to confirm this effect. Only plasmids that reproducibly conferred the pink phenotype upon reintroduction were selected for sequencing and further analysis. Transformants carrying mutations in residues previously identified as essential for BepA function—such as residue E^137^ in the active site^18^ (Extended Data Figure 1b)—were not considered further. We identified nine BepA variants of potential interest (Figure 1b), each harboring uncharacterized mutations (Extended Data Figure 1c). Collectively, these variants contained a total of 47 substitutions, including 42 unique substitutions across 40 distinct residues. These mutations are distributed throughout BepA, spanning both the TPR and M48 metalloprotease domains (Extended Data Figure 1d).

**Figure 1.**
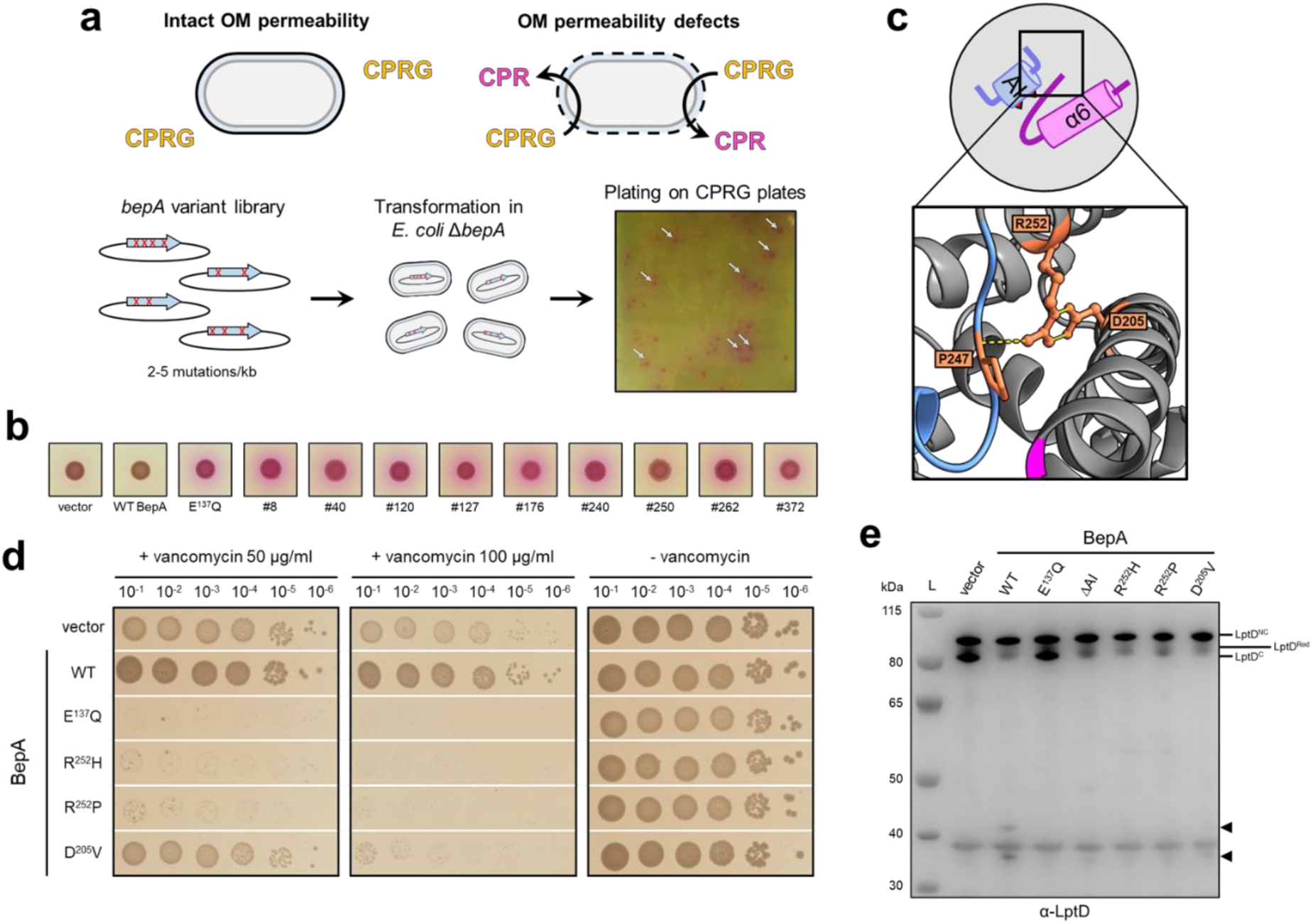
CPRG-screen reveals BepA variants inducing outer membrane defects. (**a**) Schematic illustrating the CPRG-screen. CPRG cannot penetrate the outer membrane of *E. coli*, but when outer membrane permeability defects occur, CPRG reaches the β-galactosidase LacZ, which converts it into CPR, giving a pink color to the colony. (**b**) CPRG degradation phenotype of strains producing the 9 BepA variants identified by the CPRG screen. Bacterial growth was tested in the presence of 20 µg/ml CPRG on LB agar supplemented with 0.2% L-arabinose for induction of *bepA* variants. (**c**) The side chains of D^205^ and R^252^ residues (orange) interact together via two hydrogen bonds (yellow), and the side chain of R^252^ interacts via one hydrogen bond to the backbone of the conserved P^247^ residue of the AI-plug (slate blue). (**d**) Vancomycin susceptibility phenotype of strains producing BepA with D^205^ or R^252^ substitutions. Bacterial growth was tested in the presence of 50 or 100 µg/ml vancomycin or in the absence of the drug on LB agar supplemented with 0.2% L-arabinose for induction of *bepA* variants. (**e**) *In vivo* LptD digestion assay by BepA variants. LptD was detected by Western blot in strains overproducing LptD and BepA variants. Degradation products generated by wild-type BepA are indicated by black arrows.

### Arginine 252 and aspartate 205 maintain the autoinhibitory plug in place

The most frequently substituted residue identified in the CPRG screen was R^252^, replaced by either histidine or proline in five of the nine pink clones (Extended Data Figure 1c; strains #120, #240, #250, #262, and #372). R^252^ lies within the highly conserved HPX_4_R motif shared across the M48 metalloprotease family^21^, which forms the core of BepA’s AI-plug that inserts into the active-site cleft and prevents proteolysis. In the inactive conformation, H^246^ within this motif coordinates the catalytic Zn^2+^ ion as a fourth ligand (Extended Data Figure 1b), thereby blocking access and activation of the catalytic water molecule required for peptide bond hydrolysis and locking BepA in an inhibited state^21,29^. Another residue found substituted in one of the pink clones was D^205^ (D^205^V in clone #262), a residue that is also highly conserved among BepA homologs and M48 metalloproteases (Extended Data Figure 2). Interestingly, D^205^ and R^252^ are connected by two hydrogen bonds, while R^252^ forms an additional hydrogen bond with the conserved proline residue (P^247^) of the AI-plug, suggesting that alterations in either residue may disrupt the same functional mechanism (Figure 1c). However, since mutations affecting D^205^ and R^252^ were identified alongside additional substitutions, their specific impact on BepA function needed to be determined. To this end, we tested whether BepA^R252H^, BepA^R252P^, and BepA^D205V^ could rescue the vancomycin susceptibility of Δ*bepA* cells, which results from defects in outer membrane assembly^18^. Confirming the functional importance of these residues, these mutants failed to restore resistance to vancomycin (Figure 1d). Instead, their expression exacerbated vancomycin susceptibility. This increased susceptibility could result from a dominant-negative effect previously reported for BepA catalytic dead variants such as BepA^E137Q^, which presumably bind substrates and interact with the BAM complex without completing proteolytic degradation^19,30^. Alternatively, it may result from BepA hyperactivation, causing unregulated degradation of its substrates, including the essential outer membrane protein LptD. To distinguish between these two possibilities, we assessed the impact of expressing BepA^R252H^, BepA^R252P^, and BepA^D205V^ in cells overexpressing LptD, a condition that increases the substrate load on the BAM complex and favors the accumulation of an immature form of LptD (LptD^C^), a known BepA substrate^18^. As expected, in the absence of BepA or in cells expressing the catalytically inactive BepA^E137Q^ variant, LptD^C^ accumulated (Figure 1e). In contrast, in cells expressing BepA^R252H^, BepA^R252P^, or BepA^D205V^, LptD^C^ did not accumulate; instead, the ∼35-and 40-kDa degradation products of LptD observed with wild-type BepA were absent (Figure 1e). These results suggest that mutations at D^205^ or R^252^ hyperactivate BepA, leading to complete degradation of LptD rather than the partial processing of its immature intermediate. Supporting this interpretation, a similar phenotype was observed upon expression of a hyperactive BepA variant lacking the AI-plug^21,29^ (BepA^ΔAI^, Figure 1e). Thus, D^205^ and R^252^, which are interconnected by hydrogen bonds, appear to function in tandem to stabilize the AI-plug that blocks the catalytic site, and that mutations at either residue destabilize this structure, unleashing unregulated proteolytic activity.

### The G^166^R substitution mitigates the toxic hyperactivation of BepA^R252H^

We noticed that the phenotype of strain #250, which carries the R^252^H mutation, differed from that of the other strains on CPRG: while a pinkish halo was present, the colony itself was less intensely pink (Figure 1b). This observation suggested that the three additional substitutions in this mutant (G^166^R, L^170^V, and P^239^Q), either individually or in combination, could mitigate the hyperactivation caused by the R^252^H mutation. To investigate this further, single and double mutants incorporating different combinations of the four substitutions were analyzed for vancomycin susceptibility and their effects on LptD overexpression. Among the single mutants, the L^170^V and P^239^Q substitutions had no impact on vancomycin susceptibility or LptD processing, behaving similarly to wild-type BepA (Figures 2a and 2c). In contrast, expression of BepA^G166R^, while having no effect on vancomycin susceptibility, led to the accumulation of LptD^C^ (Figure 2c). For the double mutants, the addition of L^170^V or P^239^Q to BepA^R252H^ had no effect (Figure 2b). However, introducing G^166^R in the R^252^H background fully suppressed the vancomycin susceptibility caused by R^252^H. Furthermore, whereas expression of BepA^R252H^ promotes complete degradation of LptD, cells expressing BepA^G166R/R252H^ accumulated LptD^C^ (Figure 2c). Thus, G^166^R counteracts the hyperactivation caused by R^252^H and shifts BepA toward a less active state.

**Figure 2.**
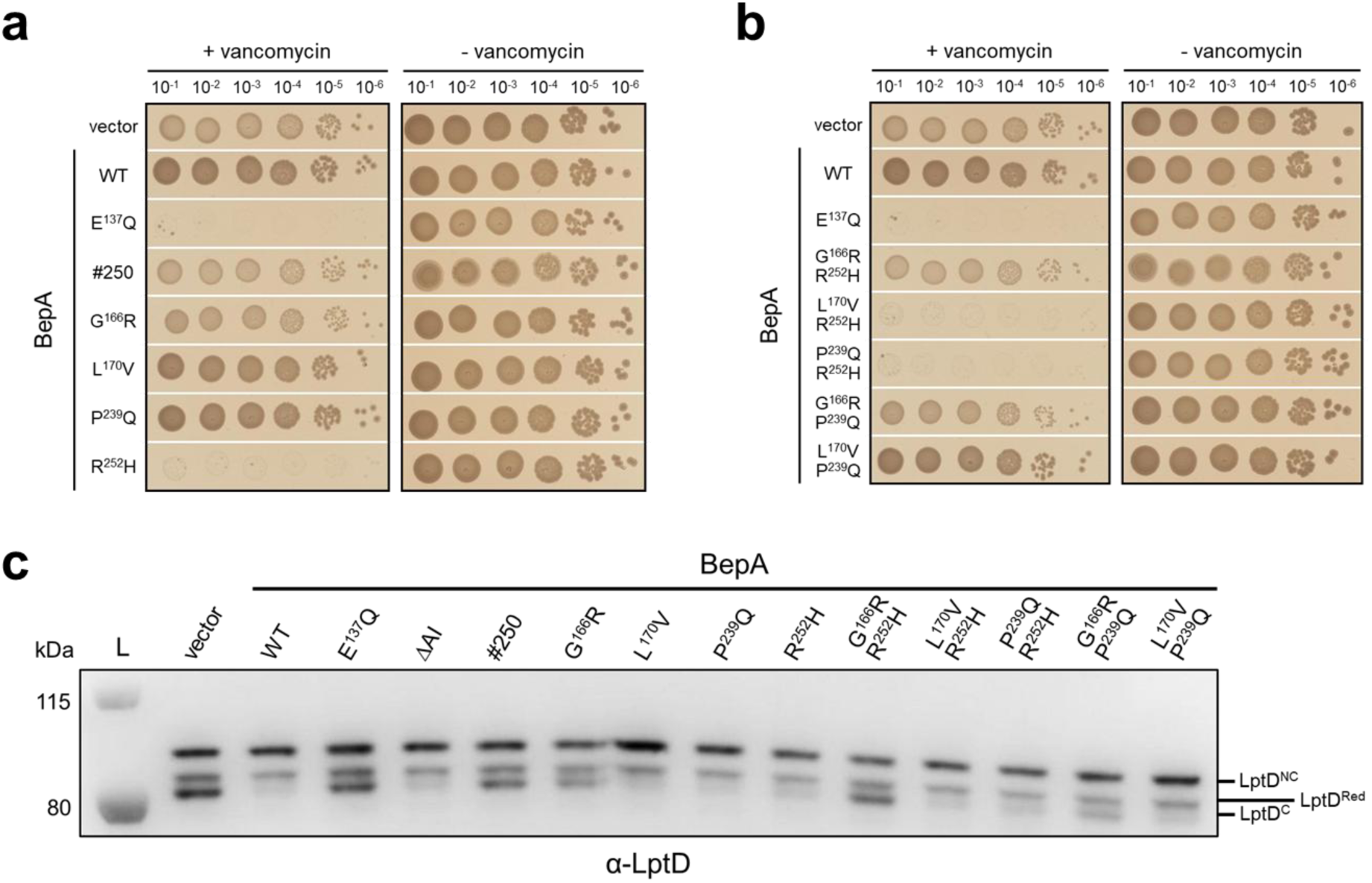
The G^166^R substitution rescues from the toxicity of BepA R^252^H. (**a**, **b**) Vancomycin susceptibility phenotype of strains producing BepA with single (**a**) or double (**b**) substitutions found in BepA #250. Bacterial growth was tested in the presence of 50 µg/ml vancomycin or in the absence of the drug on LB agar supplemented with 0.2% L-arabinose for induction of *bepA* variants. (**c**) *In vivo* LptD digestion assay by BepA variants. LptD was detected by Western blot in strains overproducing LptD and BepA variants.

### The G^166^R mutation reveals the essential role of the flexible α6-lid in BepA function

Residue G^166^ is located on an intrinsically disordered region of BepA known as the α6-lid (residues Q^154^ to T^197^), a flexible structural element^20,21^ (Extended Data Figure 1e). Interestingly, the α6-lid has been proposed to shield the substrate-binding pocket, although direct evidence for this function is still lacking (Extended Data Figures 1a and 1b). The observation that introducing the G^166^R mutation into an R^252^H background suppresses the hyperactivation phenotype suggests that mutations at G^166^ can counteract the effects of the R^252^ mutation. We hypothesized that the G^166^R substitution interferes with α6-lid dynamics, thereby restricting access to the substrate pocket and/or preventing displacement of the AI-plug. To test this hypothesis, we introduced the G^166^R mutation into the catalytic-dead mutant BepA^E137Q^. We reasoned that if access to the substrate pocket is indeed governed by α6-lid mobility, the G^166^R substitution should suppress the dominant-negative effect of E^137^Q, which normally stalls the BAM complex by binding substrates without completing proteolytic degradation. Excitingly, we found that cells expressing BepA^E137Q/G166R^ did not exhibit the severe vancomycin susceptibility observed upon expression of BepA^E137Q^ (Figure 3a). Furthermore, while BamA and LptD coeluted with BepA^E137Q^ in pulldown experiments—consistent with its proposed role in stalling the BAM complex with an incompletely folded substrate— BamA coelution with BepA^E137Q/G166R^ was markedly reduced and LptD was no longer detected (Figure 3b). Next, to determine whether physically restricting α6-lid mobility mimics the effect of G^166^R, we introduced disulfide bonds into BepA^R252H^ and BepA^E137Q^. Three cysteine pairs (Y^55^C/L^162^C, Y^55^C/L^169^C, L^60^C/L^169^C) were tested (Figure 3c). When introduced into the R^252^H or E^137^Q backgrounds, formation of these disulfides abolished vancomycin susceptibility (Figure 3d), prevented BamA and LptD from coeluting with BepA (Figure 3b), and impaired the degradation of LptD^C^ (Figure 3e), mirroring the effect of the G^166^R mutation. Altogether, these results support the idea that dynamic movement of the α6-lid is critical for BepA function, and that the G^166^R mutation perturbs this mobility, thereby impairing BepA’s engagement with the BAM complex and its substrates.

**Figure 3.**
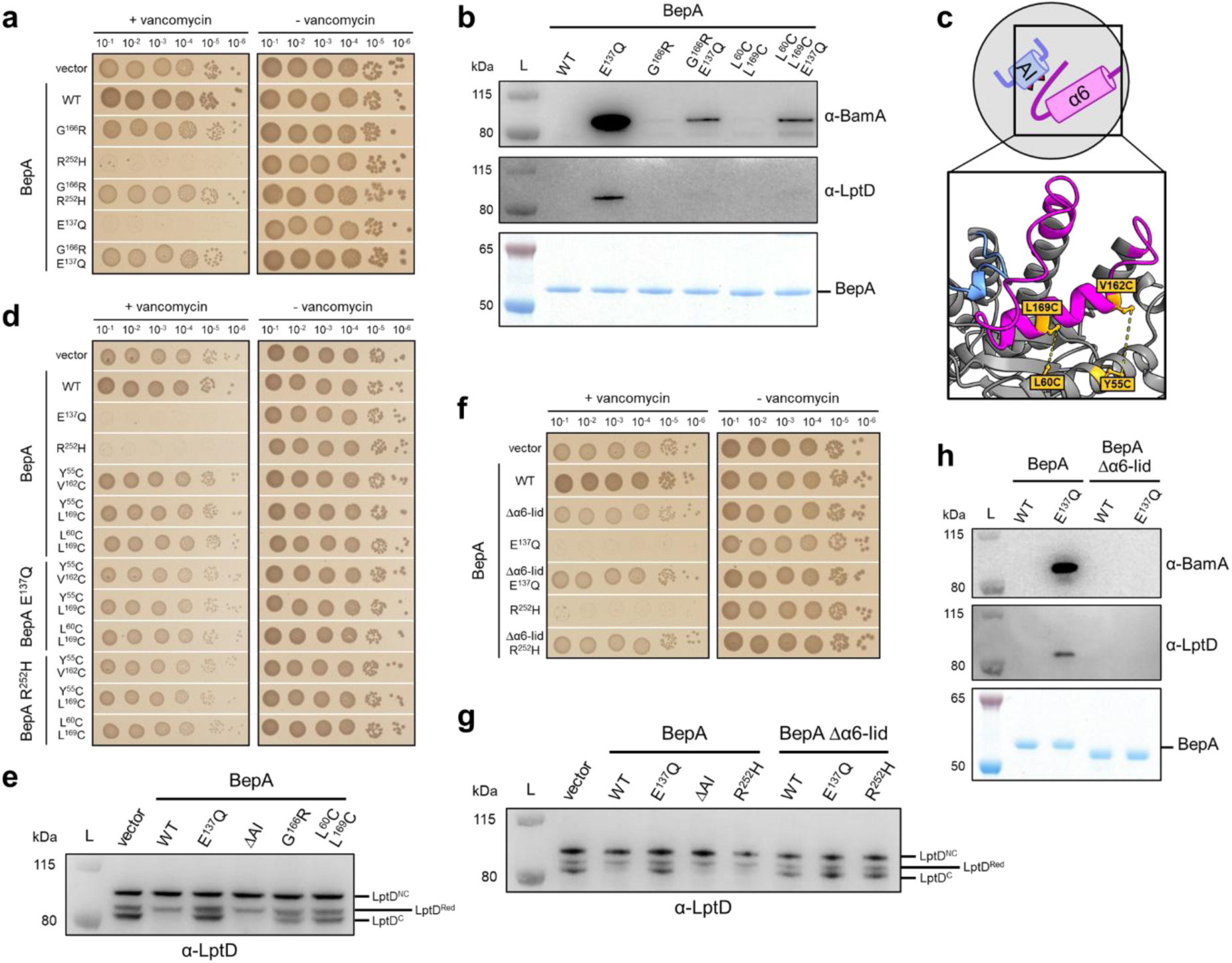
The α6-lid of BepA must open to expose its substrate pocket. (**a**) Vancomycin susceptibility phenotype of strains producing BepA or BepA G^166^R with toxic substitutions R^252^H or E^137^Q. Bacterial growth was tested in the presence of 50 µg/ml vancomycin or in the absence of the drug on LB agar supplemented with 0.2% L-arabinose for induction of *bepA* variants. (**b**) Pull-down assay on BepA variants. BepA, LptD, and BamA were detected by Western-blot after purification of strep-tagged BepA variants. (**c**) Positions of the cysteines (orange) introduced in BepA to block the α6-lid (magenta) in a closed conformation. (**d**) Vancomycin susceptibility phenotype of strains producing BepA with the α6-lid blocked in a closed conformation by disulfide bond. Bacterial growth was tested in the presence or absence of 50 µg/ml vancomycin on LB agar supplemented with 0.2% L-arabinose for induction of *bepA* variants and with 3 mM tetrathionate to increase disulfide bond formation. (**e**) *In vivo* LptD digestion assay by BepA variants. LptD was detected by Western blot in strains overproducing LptD and BepA variants. (**f**) Vancomycin susceptibility phenotype of strains producing BepA or BepA Δα6-lid with toxic substitutions E^137^Q or R^252^H. Bacterial growth was tested in the presence of 50 µg/ml vancomycin or in the absence of the drug on LB agar supplemented with 0.2% L-arabinose for induction of *bepA* variants. (**g**) *In vivo* LptD digestion assay by BepA variants. LptD was detected by Western blot in strains overproducing LptD and BepA variants. (**h**) Pull-down assay on BepA variant lacking the α6-lid. BepA, LptD, and BamA were detected by Western-blot after purification of strep-tagged BepA variants.

### The α6-lid is essential for BepA activity and interaction with the BAM complex

The results above point to the α6-lid as a critical regulatory element that governs access to the substrate pocket, interaction with BAM and its substrates, and ultimately, displacement of the AI-plug to turn BepA’s proteolytic activity on. If so, removing the α6-lid entirely should eliminate this regulatory barrier and generate a constitutively active form of BepA, potentially leading to increased degradation of LptD and exacerbation of the R^252^H phenotype. In the case of the E^137^Q variant, α6-lid deletion provides a means to assess whether exposing the substrate pocket can further increase the dominant-negative stalling effect of a catalytically inactive mutant. To test this, we complemented a Δ*bepA* strain with a plasmid carrying an allele of *bepA* in which the α6-lid was removed (deletion of residues Q^154^ to I^194^; BepA^Δα6-lid^), either alone or in combination with the catalytic-dead E^137^Q or hyperactivated R^252^H substitutions. When we deleted the α6-lid, we replaced it with a short, flexible peptide linker (GSGSGSG) to bridge the two ends of the protein, ensuring that the overall structure of BepA remained intact despite the removal of the α6-lid (Extended Data Figure 3). Unexpectedly, deletion of the α6-lid did not cause vancomycin toxicity when introduced into wild-type BepA (Figure 3f). Even more surprising, it did not increase the toxicity of the R^252^H or E^137^Q mutations; instead, it completely abolished the vancomycin susceptibility of cells expressing these mutants (Figure 3f). Moreover, deletion of the α6-lid led to a marked accumulation of LptD^C^ even in the presence of R^252^H (Figure 3g) and abrogated interactions between BepA^E137Q^ and both BamA and LptD (Figure 3h). Thus, contrary to our initial expectation that exposing the substrate pocket would enhance protease activity, deletion of the α6-lid instead suppressed BepA function—rescuing vancomycin susceptibility, preventing substrate degradation, and abolishing interactions with BAM and its clients. These unexpected findings reveal that the α6-lid not only controls access to the active-site cleft, but is also essential for BepA’s recruitment to the BAM complex and engagement with misfolded OMPs.

### Cryo-EM analysis of the BepA-BAM complex reveals association of the α6-lid with the BamA lateral gate

To gain mechanistic insight into this dual role of the α6-lid, we solved the cryo-EM structure of the BepA-BAM complex. We took advantage of the BepA #127 variant identified in the CPRG screen, which, based on pull-down experiments using strep-tagged BepA, co-purifies with higher amounts of BamA, BamB, and BamD compared to wild-type BepA (Extended Data Figure 4a). Because the catalytic-dead BepA^E137Q^ variant stalls the BAM complex (Figure 3b), we added the E^137^Q substitution to the four substitutions found in the #127 variant (A^149^G, L^169^S, Q^176^H, and G^184^C) to boost the yields of complex formation. Purification and complex stabilization were further aided by using two alternative BamA-specific His-tagged nanobodies, Nb5 and Nb32 (Supplementary Information Table S1). The BepA-BAM complexes resulting from a BepA^strep^-Nb5 or Nb32 double affinity pull-down were subjected to single particle cryo-EM and 3D reconstruction, resulting in final maps of 3.2 and 6.4 Å resolution for BepA-BAM-Nb5 and BepA-BAM-Nb32, respectively (see Methods; Extended Data Figures 4 and 5). Both complexes comprise the BAM subunits BamA, BamB, BamC, BamD and BamE, as well as BepA and the respective nanobodies, with Nb32 and Nb5 binding BamA through its extracellular L6 loop or the tip of the POTRA 1 domain, respectively (Figures 4a and 4b). In the BepA-BAM-Nb32 complex BamA is in its inward open conformation, with the β1- β16 lateral seam closed. In the BepA-BAM-Nb5 complex, however, the BamA β-barrel is found in an open, substrate-engaged conformation (Figures 4b, 4c, and 4d). The cryo-EM map shows additional density corresponding to four β-strands of an unidentified OMP substrate (possibly LptD; Figures 4b, 4c, and 4d) augmenting the BamA β-barrel at the β1 strand. In addition, the map shows an unidentified, globular density lodged inside the lumen of the BamA β-barrel (Figures 4b and 4d). In both complexes, BepA lies beneath the BamA barrel, where its TPR domain is encircled by BamD and POTRA 1, while its M48 metalloprotease domain engages POTRA 2-4, together encompassing a total contact surface area of 3,265.3 Å^2^ (Figures 4a and 4b, Extended Data Figure 6). The contacts between BepA, BamD, and POTRA 1-4 remain identical in both complexes, as does the overall structure of BepA (RMSD 1.01 Å), with the notable exception of the AI-plug (residues R^234^ to P^249^) and the α6-lid (residues Q^154^ to T^197^). In reported X-ray structures of soluble BepA (PDB ID 6AIT and 6SAR)^20,21^ the AI-plug is folded and ligates the catalytic Zn^2+^ ion through H^246^, while the α6-lid is either disordered (PDB 6SAR) or folded back against strands β1-β3 (PDB 6AIT), together rendering BepA in a catalytically locked conformation (Figure 4e). In the BepA-BAM-Nb32 complex, the absence of density for the α6-lid suggests it dissociates from the folded back position observed in 6AIT, adopting a largely disordered conformation (Figures 4e and 4g; Extended Data Figures 7d and 7e). EM density for the AI-plug region is poorly defined, indicating increased flexibility in the α8-α9-α10 loops, yet the α9 AI-plug remains coordinated to the catalytic Zn^2+^, maintaining the enzyme in a locked state. In contrast, the BepA-BAM-Nb5 complex captures the protease in an open, catalytically active conformation (Figures 4f and 4g; Extended Data Figures 7a, 7b, 7c and 7e). In this complex, opening of the BamA β-barrel and binding of the folding OMP substrate are accompanied by a ∼35° counterclockwise rotation relative to POTRA 5 (Figures 4a and 4b; Extended Data Figure 7e). This movement positions strands β15-β16 directly above the α6-lid of BepA. In turn, the otherwise disordered loop (residues P^175^ to T^197^) connecting α6 to α7 folds back and extends α7 by ∼30 Å, forming a helical hairpin with α6 that projects into the lipid bilayer and packs against F^785^, F^802^ and F^804^ of BamA β15-β16 (Figures 4b and 4f; Extended Data Figure 7c). The repositioning of the BamA β-barrel also creates a docking platform for the AI-plug, which engages the periplasmic loops connecting β13-β14 and β15-β16, further stabilized by a hydrogen bond between E^241^ and Q^191^ in the extended α7 helix. This conformational cascade results in the disengagement of His^246^ as the fourth ligand to the catalytic Zn^2+^, thereby freeing the site for water coordination and proteolysis (Figures 4f and 4g). Together, these data show that the α6-lid acts as a dynamic transmembrane element that anchors BepA to the outer membrane, positions the protease catalytic site relative to the folding substrate at the laterally opened BamA β-barrel, and forms a stabilizing docking site for the opening of the AI-plug to activate catalysis.

**Figure 4.**
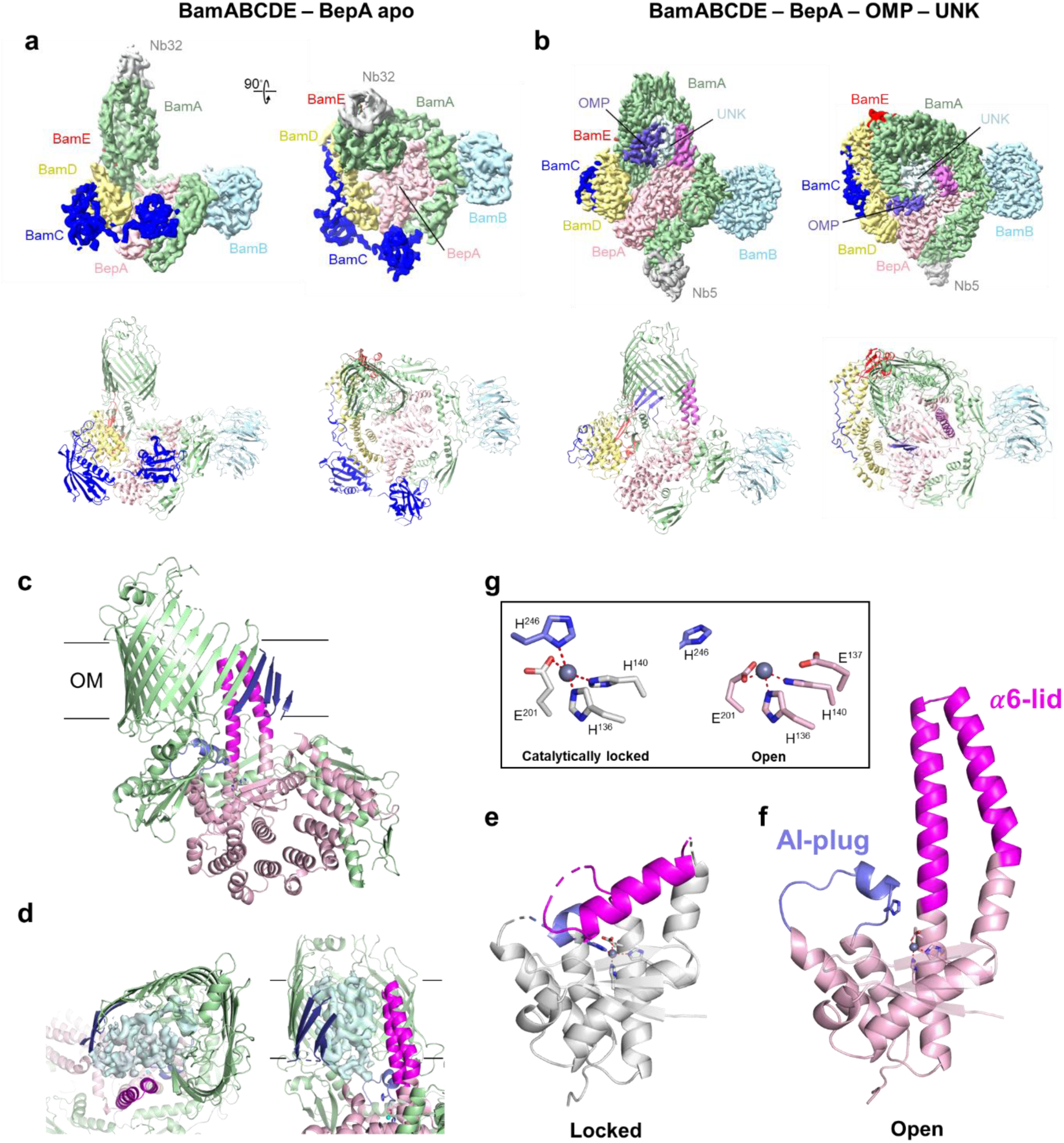
Structure of the BepA-BAM complexes. (**a, b**) Cryo-EM maps and cartoon representations of the BepA-BAM-Nb32 (**a**) and BepA-BAM-Nb5 (**b**) complexes, shown in front and top view. BamA, BamB, BamC, BamD and BamE are colored green, light blue, blue, yellow and red, respectively. BepA is shown in pink, with AI-plug and α6-lid colored slate blue and magenta, respectively. The BepA-BAM-Nb5 complex shows density for an unidentified OMP (purple) and luminal globular protein (UNK; unmodelled). (**c**) Side view of the BepA-BAM-Nb5 complex (Bam accessory subunits not shown for clarity). (**d**) Close-up cartoon representation of the BamA β-barrel, OMP substrate and BepA AI-plug and α6-lid as seen in the BepA-BAM-Nb5, including unmodelled cryo-EM density for a globular protein in the BamA β-barrel lumen. (**e, f**) Side view ribbon representation of BepA active site region (residues P^93^ to Q^269^, β1-α10) as seen in soluble, inactive BepA (PDB ID 6AIT) and the activated BepA in the BepA-BAM-Nb5 complex. (**g**) Stick representation of Zn^2+^ coordinating ligands in inactive (PDB ID 6AIT) and activated BepA (BepA-BAM-Nb5), and acid base catalyst E^137^.

### The α6-lid anchors BepA to the outer membrane under folding stress

BepA interacts with BamA via its TPR domain (Extended Data Figures 6 and 7), an association critical for its role in outer membrane protein quality control^19,20^. Our results now reveal that the α6-lid functions as a molecular harpoon, anchoring BepA to the outer membrane upon substrate stalling, a step that is essential for both its interaction with BAM and the activation of its protease activity. We next investigated whether envelope stress would serve as a molecular trigger for the membrane-anchoring mechanism that recruits BepA to the outer membrane. To explore this, we fractionated periplasmic and membrane extracts from Δ*bepA* cells expressing either BepA or BepA^Δα6-lid^. Western blot analysis revealed that both proteins are predominantly localized to the periplasm, consistent with previous reports of BepA as a soluble periplasmic protein^18^ (Figure 5a). Next, we deleted *surA*—encoding a periplasmic chaperone that escorts unfolded β-barrels across the periplasm^32–35^. Deletion of *surA* leads to the accumulation of misfolded β-barrels and activates the σ^E^ stress response. Remarkably, in the absence of SurA, BepA became exclusively membrane-associated, whereas BepA^Δα6-lid^ remained in the periplasm (Figure 5a). This observation suggests that under envelope stress—when its protease activity is most critical—BepA is recruited to the outer membrane by the α6-lid. To further probe this, we monitored GFP-tagged BepA^E137Q^ and BepA^E137Q/Δα6-lid^ by fluorescence microscopy. The E^137^Q substitution was leveraged to exacerbate the localization of BepA in the outer membrane. Cells were subjected to hyperosmotic conditions to induce plasmolysis, during which cytoplasmic shrinkage causes the periplasm to expand at the cell poles^36,37^. In Δ*bepA* cells, GFP-BepA^E137Q^ filled the expanded periplasm at the poles (Figure 5b), indicative of a mostly soluble periplasmic localization. In contrast, in Δ*bepA* cells overexpressing LptD—a condition that imposes envelope stress by burdening the assembly machinery—GFP-BepA^E137Q^ remained strongly associated with the outer membrane, while GFP-BepA^E137Q/Δα6-lid^ continued to display a soluble periplasmic distribution (Figure 5b). Together with our structural observations, these findings indicate that the α6-lid acts as a reversible transmembrane harpoon that is essential for the recruitment of BepA to the outer membrane under stress conditions.

**Figure 5.**
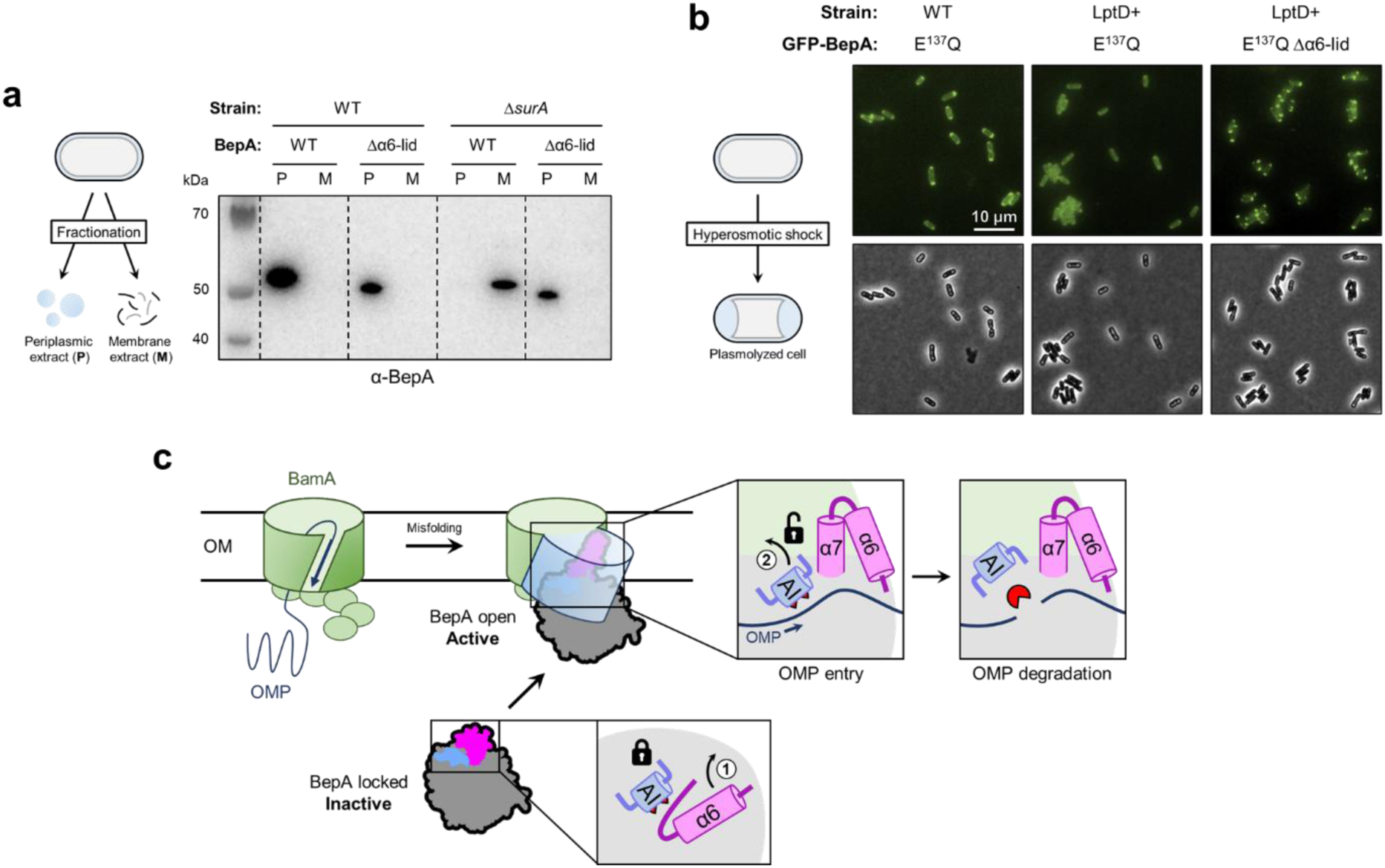
The α6-lid anchors BepA to the BAM complex in the outer membrane. (**a**) Fractionation assay to localize BepA. Periplasmic (P) and membrane (M) extracts were prepared from *E. coli* MG1655 Δ*bepA* or Δ*bepA* Δ*surA* producing BepA WT or BepA Δα6-lid. The localization of BepA within these extracts was detected by Western blot. (**b**) Microscopy assay to localize BepA. *E. coli* MG1655 Δ*bepA* or its derivative overproducing LptD (LptD+) and GFPBepA^E137Q^ or GFP-BepA^E137Q/Δα6-lid^ were grown in LB supplemented with 0.2% L-arabinose for induction of *bepA* variants and placed on an agarose PBS pad supplemented with 1 M NaCl to induce plasmolysis. The E^137^Q substitution was leveraged to exacerbate the localization of BepA in the outer membrane. (**c**) Model of BepA-mediated degradation of misfolded OMPs in the BAM complex. BepA locked-conformation lies in the periplasm as a soluble periplasmic BAM sentinel, transiently interacting with BamA through TPR-POTRA interactions. In response to misfolded OMP in BAM, the opening of the α6-lid (step 1) anchors BepA open-conformation to the BAM complex in the outer membrane. The opening of the α6-lid enables the displacement of the autoinhibitory-plug (AI; step 2) revealing the catalytic residues that degrade the misfolded OMP in the outer membrane precisely where and when it is needed.

## DISCUSSION

Because they catalyze irreversible reactions, proteases must be tightly regulated to prevent unintended damage. In Gram-negative bacteria, this is particularly true for envelope-localized proteases that ensure protein quality control in a compartment that lacks ATP and thus cannot rely on energy-dependent folding systems to remodel substrates prior to degradation. As a result, periplasmic proteases must discriminate between folding intermediates and truly defective proteins without the help of active remodeling machinery. This creates a heightened need for mechanisms that restrict protease activity to the right substrates, at the right time and place. One critical context where such control is required is the folding of OMPs by the BAM complex. This essential process is inherently error-prone, and when folding fails, misassembled OMPs stall on BAM, compromising envelope integrity and blocking access for newly arriving substrates. To maintain outer membrane homeostasis, multiple folding helpers cooperate with BAM—either by guiding nascent OMPs to the complex or by removing stalled intermediates that would otherwise clog the system^5^. Among these quality control factors, BepA plays a central role under stress conditions, ensuring the fidelity of OMP biogenesis^13,18–21,29,30^. However, its mechanism of activation remained unclear.

Here, we delineate the molecular mechanism by which BepA selectively engages the BAM complex and degrades misfolded substrates. Altogether, our results support the following model (Figure 5c): under normal conditions, BepA exists as a soluble, inactive protease in the periplasm, where its transient interactions with BamA via the TPR domain^19^ are insufficient to trigger activation. However, under envelope stress—when misfolded OMPs accumulate in BamA—an endangered substrate (such as misfolded LptD) interacts with BepA, either directly or via BamA, triggering a conformational rearrangement. This rearrangement involves the dynamic α6-lid. Upon activation, the α6-lid functions as a molecular harpoon that inserts into the outer membrane, anchoring BepA at the BAM complex. Membrane insertion not only stabilizes BepA’s association with BamA but also mechanically or allosterically dislodges the AI-plug—stabilized by hydrogen bonds network between P^247^, R^252^, and D^205^—from the catalytic residues. This displacement exposes the active site and permits water access to the catalytic Zn^2+^ center coordinated by H^246^, enabling proteolysis. Consequently, BepA can degrade misfolded substrates and restore outer membrane integrity. This sequential mechanism—initiated when a substrate misfolds on BAM and thereby recruits BepA—couples α6-lid movement, membrane anchoring, and AI-plug removal to provide precise spatial and temporal control over BepA activity. This mode of activation may represent a conserved regulatory strategy among M48 metalloproteases (see below). Whether BepA cooperates with additional factors such as the Skp chaperone—which has been shown to remove stalled substrates from BAM^14^—remains an open question. Notably, both BepA and Skp are σ^E^-regulated, suggesting a potential coordinated response to envelope stress.

A striking feature of this mechanism is the use of a flexible lid as a membrane-inserting element acting as a conditional anchor. A key question is whether this feature is conserved across the widespread M48 metalloprotease family. In addition to BepA, *E. coli* encodes three other M48 metalloproteases: the outer membrane lipoproteins LoiP and YcaL, and the integral inner membrane protein HtpX. AlphaFold3 predicts that all three homologs retain a hydrophobic lid homologous to the α6-lid of BepA, yet in all cases, the lid adopts an open conformation (Extended Data Figure 8a). A similar lid in the same conformation is also observed in the endoplasmic reticulum protease Ste24/ZMPSTE24 (in yeast and mammals) and in the mitochondrial inner membrane protease OMA1^23–28^ (Extended Data Figure 8b). Unlike BepA, however, these proteases are already membrane-associated—either through lipidation, as in LoiP and YcaL, or via transmembrane segments, as in HtpX, Ste24/ZMPSTE24, and OMA1. In this context, the lid may primarily act as a gate controlling substrate access rather than as a membrane anchor. Nevertheless, in HtpX, Ste24/ZMPSTE24, and OMA1, the lid lies adjacent to transmembrane helices, raising the possibility that it may transiently interact with the membrane or modulate membrane engagement in response to stress. Since all known M48 metalloproteases are membrane-associated—with BepA being the only soluble example—we propose that BepA evolved from a membrane-bound ancestor that lost its transmembrane helices and redeployed the α6-lid as a flexible docking device.

Notably, the triggerable harpoon mechanism we identified in BepA may extend beyond the M48 family. AsmA-like phospholipid transporters such as YhdP and YhjG also carry conserved hydrophobic α6-lid-like loops predicted to integrate into the outer membrane^38,39^ (Extended Data Figure 8c). These features may reflect a broader harpooning strategy by which periplasmic or membrane-associated proteins reversibly dock in response to stress signals. In BepA, this mechanism ensures tight spatiotemporal control of protease activity; in other systems, it may regulate transport, signaling, or membrane contact site formation. This new mechanism of periplasmic protein anchoring to the outer membrane—which could be help by dedicated OMPs—may represent a widespread and previously overlooked strategy among diderm bacteria.

Our work also unveiled two key residues that stabilize the AI-plug and downregulate BepA activity. These residues, D^205^ and R^252^, form a network of hydrogen bonds with the conserved P^247^ residue carried by the plug (Figure 1, Extended Data Figures 1 and 2). Together, they form a regulatory triad that acts as a safety switch, keeping the protease inactive. This AI-plug-based control mechanism is conserved across M48 metalloproteases and may be exploitable for therapeutic intervention. In *Staphylococcus aureus*, for instance, the β-lactam sensor BlaR1 remains inactive until it binds a β-lactam antibiotic with its sensor domain. This event activates the M48 metalloprotease domain—by displacing the plug—triggering cleavage of the BlaI repressor and inducing production of the β-lactamase BlaZ and the low-affinity transpeptidase MecA^40,41^. Inhibiting BlaR1 protease activity could block this resistance pathway and provide a route to combat methicillin-resistant *S. aureus*. Similar strategies may apply to the human M48 metalloproteases OMA1 and ZMPSTE24. Similar strategies may extend to the human M48 metalloproteases OMA1 and ZMPSTE24. OMA1 is an attractive drug target in diseases ranging from heart failure to cancer due to its role in mitochondrial dysfunction and cell death^42–44^. For ZMPSTE24, activating the protease could counteract loss-of-function mutations that cause premature aging^45^ and may boost antiviral defense, consistent with its role in restriction of SARS-CoV-2 and other enveloped viruses^46,47^.

In conclusion, our work reveals a membrane-coupled activation mechanism that ensures precise control of BepA’s proteolytic function. By integrating substrate recognition, membrane docking, and autoinhibitory release into a single coordinated process, this system minimizes off-target degradation while rapidly clearing toxic intermediates. The underlying principles—harpoon-like insertion, lid dynamics, and plug regulation—may be broadly conserved across the M48 metalloprotease family, in bacteria and beyond.

## METHODS

### Strains, plasmids, and growth conditions

All strains were derived from *E. coli* MG1655. The origin and genotype of strains, and the origin and characteristics of plasmids are listed in Supplementary Table S1. Bacteria were grown in Miller’s Lysogeny Broth (LB; Difco) broth or agar at 37 °C unless otherwise specified. Bacterial liquid cultures were aerated by vigorous shaking (200 rpm). P1 transduction^51^ of the Km^R^ cassette from selected mutants of the Keio collection^48^ was used to introduce gene deletions in *E. coli* MG1655. The growth media were systematically supplemented with drugs to counter-select plasmid loss (Supplementary Table S1). Spectinomycin and ampicillin were both used at 100 µg/ml. Kanamycin at 50 µg/ml was used for the Km^R^ cassette. Induction of *araBAD* promoter was performed with L-arabinose at 0.2%. Plasmids constructed in this study were obtained by using NEBuilder HiFi DNA assembly (New England Biolabs) method, unless otherwise specified. All plasmids used for the expression of *bepA* variants derived from the pBAD42-*bepA* plasmid.

### Generation of *bepA* variant plasmid library

Random, unbiased mutations in *bepA* were introduced using the GeneMorph II EZClone Domain Mutagenesis Kit (Agilent), with an expected mutation rate of 2-5 mutations per kb. Reactions were performed according to the manufacturer’s instructions with minor modifications. In the first PCR, *bepA* was amplified from 250 ng of pBAD42-*bepA* with the kit’s DNA polymerase to introduce random mutations. The PCR product was digested with DpnI, purified, and 500 ng was used as a megaprimer for a second PCR with Q5 DNA polymerase on 50 ng of pBAD42-*bepA*. The resulting PCR product was purified and transformed into NEB 5-alpha competent cells. Transformants were plated on LB agar containing 100 µg/ml spectinomycin and incubated for 16 h at 37 °C. Serial dilutions were plated in parallel to estimate library size. After growth, colonies were scraped, pooled, and stored as aliquots at-80 °C. The *bepA* variant plasmid library was purified from one aliquot by miniprep extraction.

### CPRG-based screening for outer membrane permeability defects

The *bepA* variant plasmid library was transformed into *E. coli* MG1655 Δ*bepA*. The transformants were diluted, plated on LB agar supplemented with 100 µg/ml spectinomycin (to counter-select plasmid loss), 0.2% L-arabinose (to induce *bepA* expression), and 20 µg/ml chlorophenol red-β-D- galactopyranoside (CPRG; to identify BepA-induced outer membrane defects), and grown for 16 h at 37 °C. Colonies displaying a pink phenotype were streaked onto fresh LB agar with the same supplements to confirm the phenotype. To exclude chromosomal mutations as the cause of the phenotype, plasmids were extracted from pink colonies and retransformed into a fresh *E. coli* MG1655 Δ*bepA* background. Transformants were again plated on selective CPRG-containing medium. Plasmids that reproducibly induced outer membrane permeability defects were sequenced to identify mutations in *bepA*.

### Plating efficiency assay

Bacteria were grown at 37 °C with vigorous shaking to the late growth phase, *i.e.* to an OD_600nm_ 1.0 to 4.0. The OD_600nm_ was adjusted to 1.0 and 10-fold serial dilutions (10^−1^ to 10^−6^) were prepared in LB broth. Of the resulting bacterial suspensions, 5 µl were spotted on LB agar supplemented with inducers and drugs as indicated in the legend to figures. For the CPRG phenotypic assay, 5 µl of the bacterial suspension at an OD_600nm_ of 0.1 was spotted on LB agar supplemented with inducers and drugs as indicated in the legend to figures. Plates were imaged after 16 h of incubation at 37 °C. Data shown in the figures are representatives of at least three biological repeats.

### Immunoblotting

Bacteria were grown at 37 °C to the early exponential growth phase, *i.e.* to an OD_600nm_ 0.05 to 0.1. To induce expression of *bepA* alleles, L-arabinose was added to a final concentration of 0.2% and the bacteria were further grown for 1 h at 37 °C to the late exponential growth phase, *i.e.* to an OD_600nm_ 0.4 to 0.8. The OD_600nm_ was monitored and 900 µl of the bacterial culture were precipitated on ice in 100 µl of a 100% trichloroacetic acid solution. The precipitate was centrifuged and washed with 1 ml of ice-cold acetone. Acid-denatured proteins were solubilized in SDS-sample buffer (60 mM Tris-HCl pH 7.4, 2% SDS, 10% glycerol, 0.1 mg/mL bromophenol blue). The amount of protein was normalized between the samples by adding 100 µl of SDS-sample buffer for 1 OD_600nm_ unit. Samples were boiled for 10 min before being loaded onto a precast NuPAGE Bis-Tris 4-12% gel (Life Technologies). Immunoblotting was performed according to standard procedures using 1:5,000 anti-LptD antibody (rabbit serum, CER group, Belgium), 1:10,000 anti-BepA antibody (gift from Yoshinori Akiyama), 1:30,000 anti-BamA antibody (gift from Thomas J. Silhavy), 1:10,000 anti-BamB antibody (rabbit serum, CER group, Belgium), or 1:10,000 anti-BamD antibody (rabbit serum, CER group, Belgium) followed by a horseradish peroxidase-conjugated anti-rabbit antibody (Sigma). For visualization of LptD degradation products, immunoblotting was performed with 1:50,000 anti-LptD antibody from Yoshinori Akiyama^29^. Chemiluminescence was performed with Immobilon Western Chemiluminescent HRP Substrates (Millipore) and imaged on an Amersham ImageQuant 800 Western blot imaging system (Cytiva).

### Pull-down assay

The derivatives of pBAD42 encoding BepA variants with a strep-tag between the A^27^ and D^28^ residues were transformed in MG1655 Δ*bepA*. Bacteria were grown at 37 °C to the late exponential growth phase, *i.e.* to an OD_600nm_ 0.4 to 0.6. To induce expression of *bepA* alleles, L-arabinose was added to a final concentration of 0.2% and the bacteria were further grown for 3 h at 37 °C. Cells were pelleted, resuspended in lysis buffer (50 mM Tris-HCl pH 8, 300 mM NaCl, 1 mM MgCl_2_, 1% DDM, 1 µg/ml DNase I, 50 µg/ml lysozyme, and cOmplete EDTA-free protease inhibitor cocktail (Roche)), and lysed by slight agitation on a tube-roller for 2 h at 4 °C. The lysate was clarified by centrifugation and BepA variants were purified from the clarified lysate by streptavidin affinity chromatography 20 mM Tris-HCl pH 8, 150 mM NaCl, supplemented with 2.5 mM D-desthiobiotin for the elution. SDS-sample buffer was added to the elution, and the samples were boiled for 10 min before being loaded onto a precast NuPAGE Bis-Tris 4-12% gel (Life Technologies). The volume of loaded samples was adjusted to load the same amount of BepA for each sample. Immunoblotting was performed according to standard procedures.

### Generation, Selection, and purification of nanobodies against BamA

The *E. coli* protein BamA was produced and purified as described previously^52^. An alpaca was immunized with six injections of 0.1 mg of the recombinant BamA (Tris 20 mM pH 8.0, NaCl 150 mM, 0.03 % DDM) over a six-week period. Blood was collected 4 days after the last immunization for the construction of the immune library and nanobody selection using a phage display protocol^53^. In brief, lymphocytes were isolated from the collected blood, following by the total RNA purification and cDNA synthesis using reverse transcription. The cDNA was used as template for PCR amplification of sequences corresponding to the variable domains of the heavy chain only antibodies and cloned into a pMESy4 phagemid vector (GenBank KF415192) to create the nanobody library. After three rounds of panning, 37 BamA binders were selected that belong to 35 unique families based on their sequence variation in CDR1, 2 and 3. Epitope mapping showed the selected nanobodies corresponded to seven groups with non-overlapping epitopes, two of which extracellular, 5 targeting the BamA POTRA domains. Two nanobodies, SVN_Nb5 and SVN_Nb32 (Supplementary Table S1), were selected for structural studies, binding periplasmic and extracellular epitopes, respectively. The BamA nanobody derivatives in pMESy4 were transformed into *E. coli* WK6 for overexpression. The WK6 cells were grown in Terrific Broth medium containing 0.1 % glucose and 100 µg/mL ampicillin to an OD_600nm_ of 0.8, induced with 1 mM IPTG, and incubated at 28 °C for 16 h. The periplasmic extraction was done by osmotic shock. Cells were resuspended in 20% w/v sucrose, 100 mM Tris-HCl pH 8.0, 1 mM EDTA, spun down and resuspended in ice-cold water for the release of periplasmic content by osmotic shock. His-tagged nanobodies were purified from the periplasmic extract using Ni-NTA immobilized metal affinity chromatography. Purified nanobodies were concentrated (Amicon-Ultra 10-kDa cut-off) and loaded on a HiLoad 10/30 Superdex 75 gel filtration column equilibrated in phosphate buffer saline (PBS) for further purification. The epitope mapping of BamA binders was done on full-length BamA and a POTRA-only BamA truncate, via Biolayer interferometry using an OctetRed96-FortBio with black 96-well plates-Greiner. Briefly, purified full-length and POTRA BamA (the BamA antigens) were first biotinylated using the EZ-Link NHS-PEG4-biotin kit (Perbio Science). The BamA antigens or His-tagged nanobodies were immobilized on streptavidin or Ni-NTA biosensor tips, followed by a wash in binding buffer (PBS containing 0.03 % DDM and 5 mM Imidazole). The binding (association and dissociation) of the BamA antigens and nanobodies were monitored for several minutes by dipping the tips in various concentrations of binders or BamA antigens. Nanobodies were grouped into epitope overlap groups based on mutually exclusive binding to BamA constructs, monitored by consecutively dipping of the sensor tips into different nanobody solutions.

### Purification of the BepA-BAM-Nb complexes for cryoEM

*E. coli* BW25113 competent cells were transformed with the pBAD42-*bepA*^#127/E137Q^ plasmid, containing strep-tagged BepA variant #127 (A^149^G, L^169^S, Q^176^H, and G^184^C) with additional catalytic site mutation E^137^Q. An overnight pre-culture was used to inoculate LB broth containing 50 µg/ml spectinomycin, and the culture was grown at 37 °C to an OD_600nm_ of 0.8. BepA overexpression was induced by adding 0.2% L-arabinose at 37 °C for 3 h. Cells were harvested and membrane protein were extracted by a detergent induced whole cell lysis protocol as previously described^52^. Briefly, cell pellets were resuspended in TBS buffer (50 mM Tris pH 8.0, 300 mM NaCl), supplemented with 100 μg/ml lysozyme, 50 μg/ml DNAse, 10 mM MgCl_2_, 0.4 mM AEBSF, and 1 µg/mL leupeptin, to a final OD_600nm_ of 100. DDM was added to the mixture to obtain a final concentration of 1.5% DDM. Cell lysis was done at 4 °C for 2 hours with gentle shaking. The lysates were clarified using an ultracentrifuge with 45 Ti rotor (Beckman) at 40,000 RPM and 4 °C for 45 min. BepA^#127/E137Q-strep^ was purified using Strep-Tactin®XT 4Flow® high-capacity resin (IBA), following the manufacturer protocol, and eluted in TBS buffer supplemented with 0.05% DDM, 0.02% LMNG and 50 mM biotin (IBA). To separate BAM-engaged BepA from the pool of free periplasmic BepA, the BepA^#127/E137Q-strep^ elution was mixed with BamA-binding nanobodies Nb32 or Nb5, and subjected to a second affinity purification using the Nb C-terminal hexa-histidine tag. To do so, the mixture was then loaded onto an equilibrated 0.2 ml HisPur™ Ni-NTA Spin Columns (Thermo-Fisher). Unbound proteins were washed with TBS supplemented with 0.01% LMNG and 50 mM imidazole. BepA-BAM-Nb32/Nb5 complexes were then eluted with TBS supplemented with 0.01% LMNG and 300 mM imidazole, analyzed with gel filtration, semi-native SDS PAGE, Western blot analysis, and electron microscope. This double pull-down procedure results in the isolated of recombinant BepA^#127/E137Q-strep^ engaged with the native BAM complex, as well as OMP substrates in the case of Nb5 (Extended Data Figures 4 and 5).

### Cryogenic Electron Microscope sample preparation and data collection

For graphene oxide grid preparation, 3 µl of 0.2 mg/ml graphene oxide (GO, Sigma-Aldrich) was spotted on glow-discharged R2/1 copper 300 mesh grids (Quantifoil) and incubated for 2 minutes before blotting the excessive GO. The grids were rinsed once with water, blotted and dried for at least 20 min before use. For sample application, 3 µl purified BepA-BAM-Nb complexes (∼0.1 mg/mL) were applied onto GO grids, subjected to 4 s backside blotted to remove excess liquid, and flash-frozen in liquid ethane using a CP3 plunger (Gatan). Grids were then stored in liquid nitrogen or loaded into a 300-kV CRYO ARM 300 (JEM-Z300FSC) Field Emission Cryo-Electron Microscope (JEOL) equipped with a K3 Summit direct electron detector (Gatan). Image stacks were recorded in counting mode with a cumulative electron dose of 60 electrons per Å^2^ spread over 60 frames, at 60K magnification, using serialEM^54^. Multi-hole shots were deployed to accelerate data collection.

### Cryogenic Electron Microscope data processing and 3D reconstruction

MotionCor1.3 was used to correct the frame motion^55^ and the defocus values were estimated by CtfFind4^56^ with Dose-weighting scheme. Particles were picked by crYOLO using a pre-trained model^57^. Particle extraction was done in Relion^58^ and exported to cryoSPARC for several 2D classification to discard the junk particles^59^. Classes containing the different BepA-BAM complexes were then imported back to Relion for re-extraction, ab-initio 3D reconstruction, and 3D classification with BLUSH. The final classes were used for 3D refinement with BLUSH or imported to cryoSPARC for 3D non-uniform refinement.

### Model building, refinement and validation

To make the initial model, the structure of BAM open conformation (PDB ID 7YE4), BAM closed conformation (PDB ID 9CNW), and BepA (PDB ID 6AIT) was docked into the refined map of BepA-BAM using UCSF Chimera^60^. The initial model was then manually built/edited using COOT^61^. Final model was refined using PHENIX real-space refinement^62^. Structure and map display and figures were prepared using UCSF Chimera, UCSF ChimeraX^63^, and PyMOL (https://pymol.org/2/). Data collection and refinement statistics are found in Supplementary Table S2.

### Cell envelope fractionation

Bacteria were grown at 37 °C to the early exponential growth phase, *i.e.* to an OD_600nm_ 0.05 to 0.1. To induce expression of *bepA* alleles, L-arabinose was added to a final concentration of 0.2% and the bacteria were further grown for 1 h at 37 °C to the late exponential growth phase, *i.e.* to an OD_600nm_ 0.4 to 0.8. Cells were centrifuged, resuspended on ice in 100 mM Tris-HCl pH 8.0, 20% D-sucrose, 1 mM EDTA, and incubated for 15 min on a tube-roller at 4 °C. Cells were centrifuged, resuspended on ice in 5 mM MgSO_4_, and incubated for 15 min on a tube-roller at 4 °C. Cells were centrifuged and the supernatant corresponding to the periplasmic extract was collected. The cell pellet was resuspended in 25 mM Tris-HCl pH 8.0, 1 mM EDTA and disrupted by sonication. The lysate was ultracentrifuged at 100,000 g for 1 h at 4 °C, the pellet corresponding to the membrane extract was resuspended in 25 mM Tris-HCl pH 8.0 supplemented with 1% DDM and incubated overnight on a tube-roller at 4 °C. Protein concentrations of periplasmic and membrane extracts were determined with the Microplate BCA Protein Assay Kit (Thermo Scientific). SDS-sample buffer was added to normalized protein quantity and the samples were boiled for 10 min before being loaded onto a precast NuPAGE Bis-Tris 12% gel (Life Technologies). Immunoblotting was performed according to standard procedures.

### Fluorescence microscopy imaging

Bacteria were grown at 37 °C to the early exponential growth phase, *i.e.* to an OD_600nm_ 0.05 to 0.1. To induce expression of *bepA* alleles, L-arabinose was added to a final concentration of 0.2% and the bacteria were further grown for 1 h at 37 °C to the late exponential growth phase, *i.e.* to an OD_600nm_ 0.4 to 0.8. Cells were loaded on a 1% agarose PBS pad supplemented with 1 M NaCl to induce plasmolysis. Cells were imaged on a Nikon Ti2-E fully motorized inverted epifluorescence microscope equipped with a CFI Plan Apochromat λ DM 100×1.45/0.13 mm Ph3 oil objective (Nikon), a Sola SEII FISH illuminator (Lumencor), and a Prime95B camera (Photometrics). In all experiments, the msfGFPox variant of GFP was used to enhance the fluorescence signal in the periplasm.

## DATA AVAILABILITY

Data that support the findings of this study are available in the Article. Source data are provided with this paper. Cryo-EM potential maps and 3D coordinates of the reported complexes have been deposited at the EMDB and PDB under the following accession codes: BepA-BAM-Nb32 (EMD-55064; 9SOX); BepA-BAM-Nb5 (EMD-55063; 9SOW).

## ACKNOWLEDGMENTS

We thank Y. Akiyama for his kind gift of anti-LptD and anti-BepA antibodies. This work was supported by the Fonds de la Recherche Scientifique (FRS-FNRS) (grant WELBIO-CR-2022 to J.-F.C). H.V. was supported by a post-doctoral EMBO fellowship (ALTF 359-2023). We acknowledge financial support from VIB, FWO EOS (grant G0G0818N to H.R.) and FWO research infrastructure (grant G0H5916N to H.R.). S.-V.N. was partially supported by a senior post-doctoral FWO fellowship (12ZM421N).

## AUTHOR CONTRIBUTIONS

H.V., P.C.N., and S.-V.N performed the experiments. H.V., P.C.N., S.-V.N, P.L., B.I., S.-H.C., H.R., and J.-F.C. analyzed the data. S.-H.C. and J.-F.C. conceived the project and designed the study. H.V., H.R., and J.-F.C. wrote the paper with contributions from all authors.

---

**Extended Data Figure 1.**
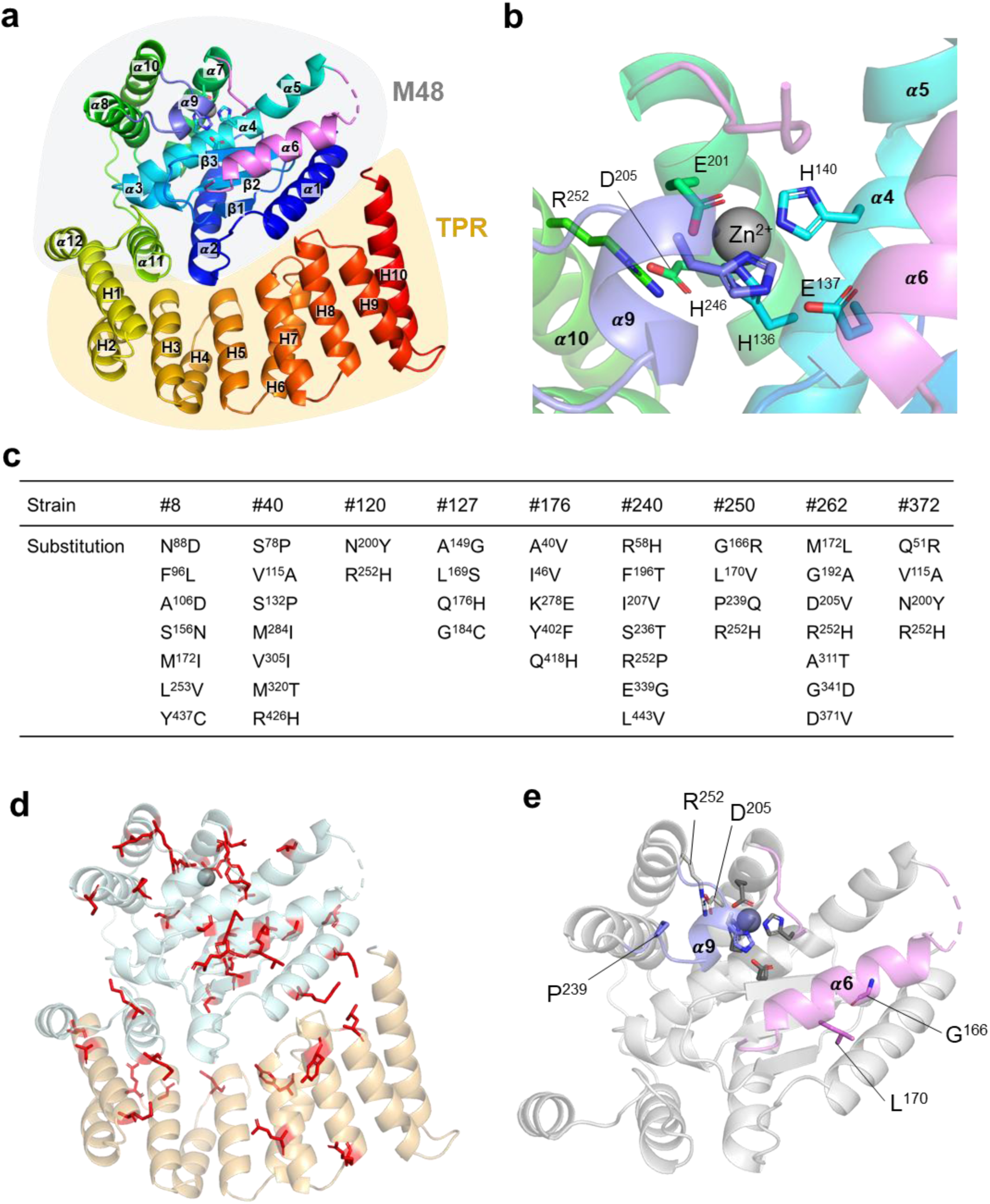
BepA structure and variants. (**a**) Structure of *E. coli* BepA (PDB ID 6AIT) shown in cartoon representation and colored blue to red from N-to C-terminus. The α9 autoinhibitory loop (AI-plug) and α6-lid are colored slate blue and magenta, respectively. Secondary structure elements in the catalytic M48 metalloprotease domain are labeled α1-α12 and β1-β3, those in the C-terminal tetratricopeptide repeat (TPR) domain are labeled H1-H10. (**b**) Close-up view of the BepA catalytic side, with the active site residues and catalytic Zn^2+^ ion shown as sticks and sphere, respectively. The Zn^2+^ ion is coordinated by H^136^, H^140^ and E^201^ in α4 and α7. The fourth ligation site is coordinated by H^246^ in the AI-plug, thereby blocking water activation and catalysis. The catalytic site is completed by E^137^, acting as essential acid/base catalyst. D^205^ and R^252^ stabilize the AI-plug. Colored as panel a. (**c**) List of the 47 substitutions identified, comprising 42 unique substitutions across 40 different residues. (**d**) Localization of the substitutions scattered throughout BepA’s structure. Substituted residues are in stick representation and colored red. (**e**) Close-up of BepA M48 metalloprotease domain with D^205^ and residues in the #250 mutant highlighted in stick representation. The AI-plug (containing α9) and α6-lid are colored slate blue and magenta, respectively.

**Extended Data Figure 2.**
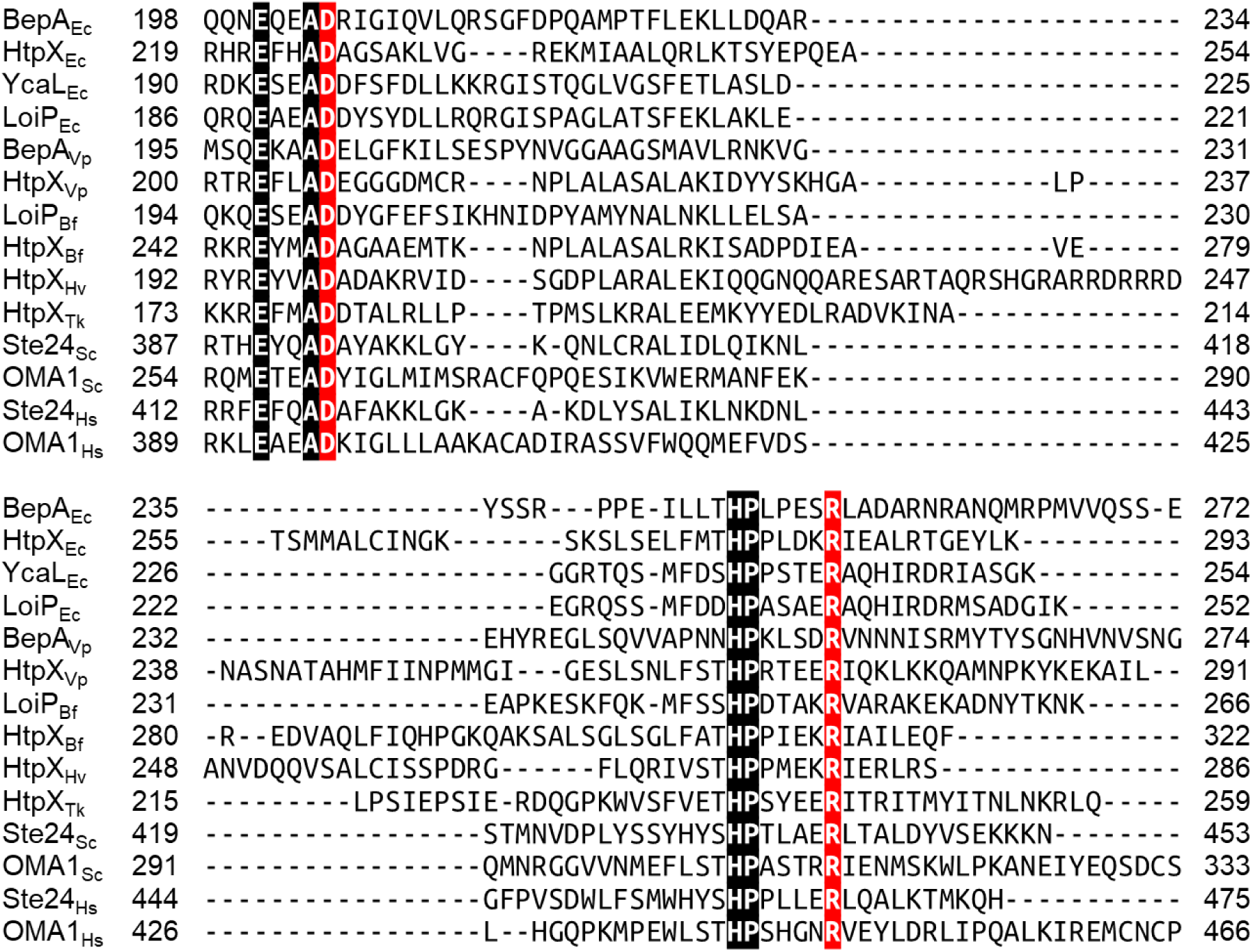
Sequence alignment of M48 metalloproteases. For clarity, only the regions containing BepA’s D^205^ and R^252^ residues (highlighted in red) are shown. Other conserved residues are highlighted in black. The alignment was generated by Clustal Omega Multiple Sequence Alignment. Ec, *Escherichia coli*; Vp, *Veillonella parvula*; Bf, *Bacteroides fragilis*; Hv, *Haloferax volcanii*; Tk, *Thermococcus kodakarensis*; Sc, *Saccharomyces cerevisiae*; Hs, *Homo sapiens*.

**Extended Data Figure 3.**
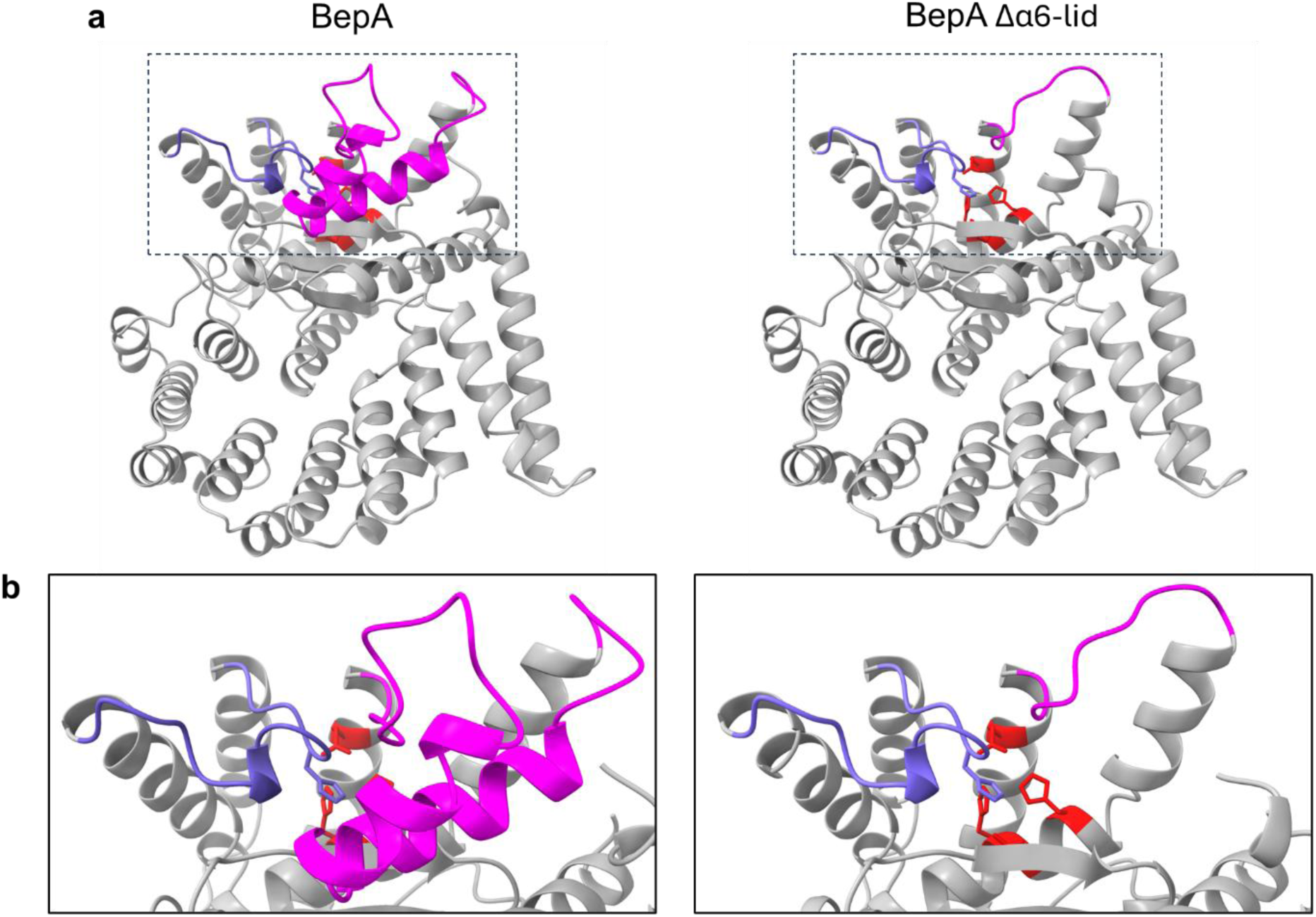
Structural comparison of wild-type BepA and BepA Δα6-lid replaced by a GSGSGSG linker. (**a**) AlphaFold3 predictions of BepA structures shown in cartoon representation. (**b**) Boxed area showing the catalytic site. BepA is colored gray, with the AI-plug and α6-lid colored slate blue and magenta, respectively. Catalytic residues are colored red. Zn^2+^ ligands are shown in stick representation.

**Extended Data Figure 4.**
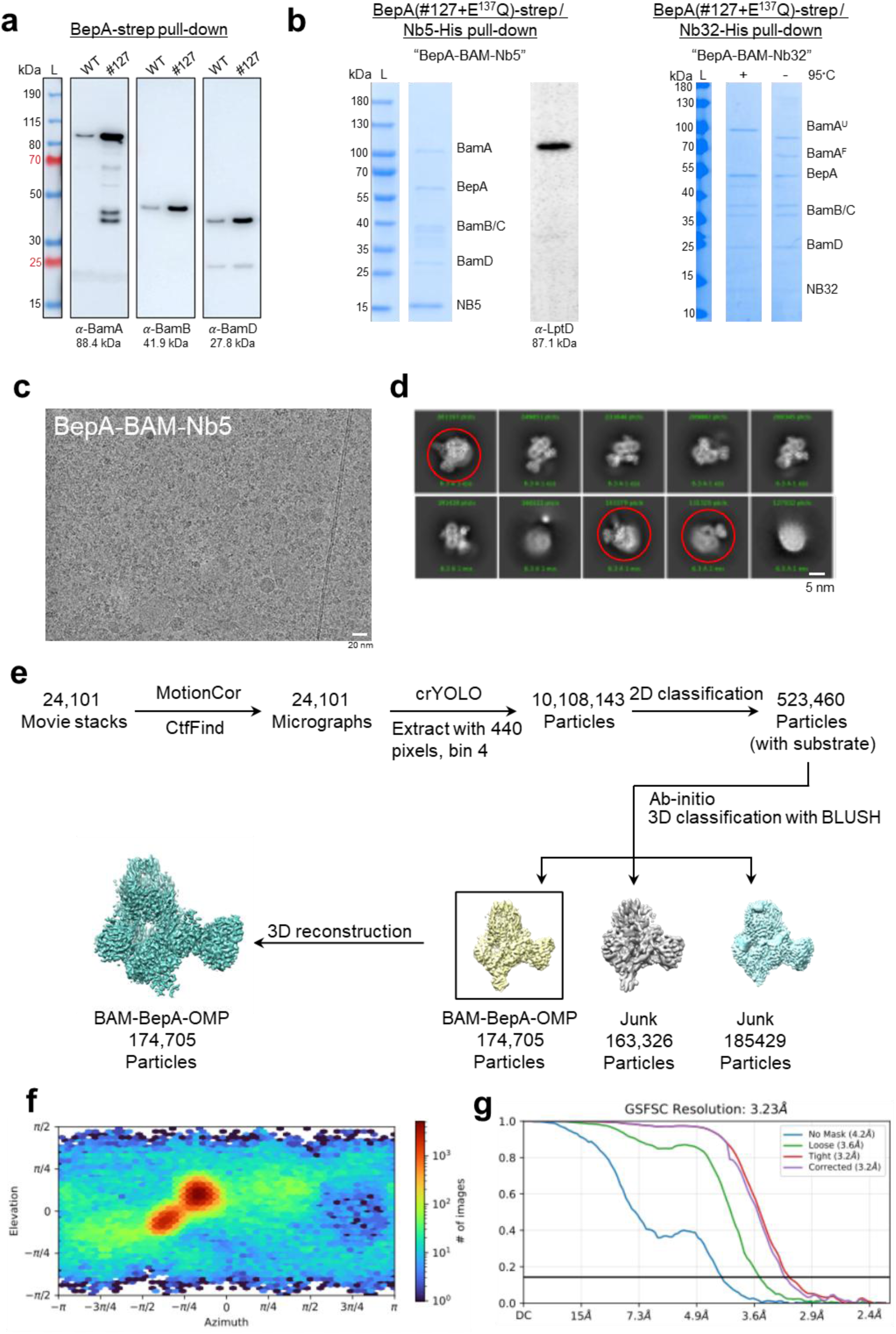
BepA complex purification and cryoEM 3D reconstruction of BepA-BAM-Nb5 complex. (**a**) BepA #127 copurifies with more BAM complex. Pull-down assay on BepA WT and BepA #127. The amount of BepA, BamA, BamB, and BamD were detected by Western-blot after purification of strep-tagged BepA variants. (**b**) SDS PAGE and western blot (anti-LptD) of BepA-BAM-Nb5 and BepA-BAM-Nb32 complexes used for cryoEM analysis. Fractions shown are the elution of the affinity pulldown by SVN_Nb5 and SVN_Nb32, respectively. The pull-down sample includes LptD, though it may contain additional OMPs. (**b**) Raw cryoEM micrograph of BepA-BAM-Nb5 collected at 300kV and 60k magnification. (**c**) 2D classes obtained for BepA-BAM-Nb5 complex. (**d**) Image processing and structure determination pipeline used for the BepA-BAM-Nb5 dataset, resulting in the 3D reconstruction of a complex containing BamABCDE-BepA-Nb5 and an unidentified OMP substrate. (**e, f**) angular distribution map of the particles used (e), and FSC curves (f) of the final reconstruction of the BAM-BepA-OMP complex.

**Extended Data Figure 5.**
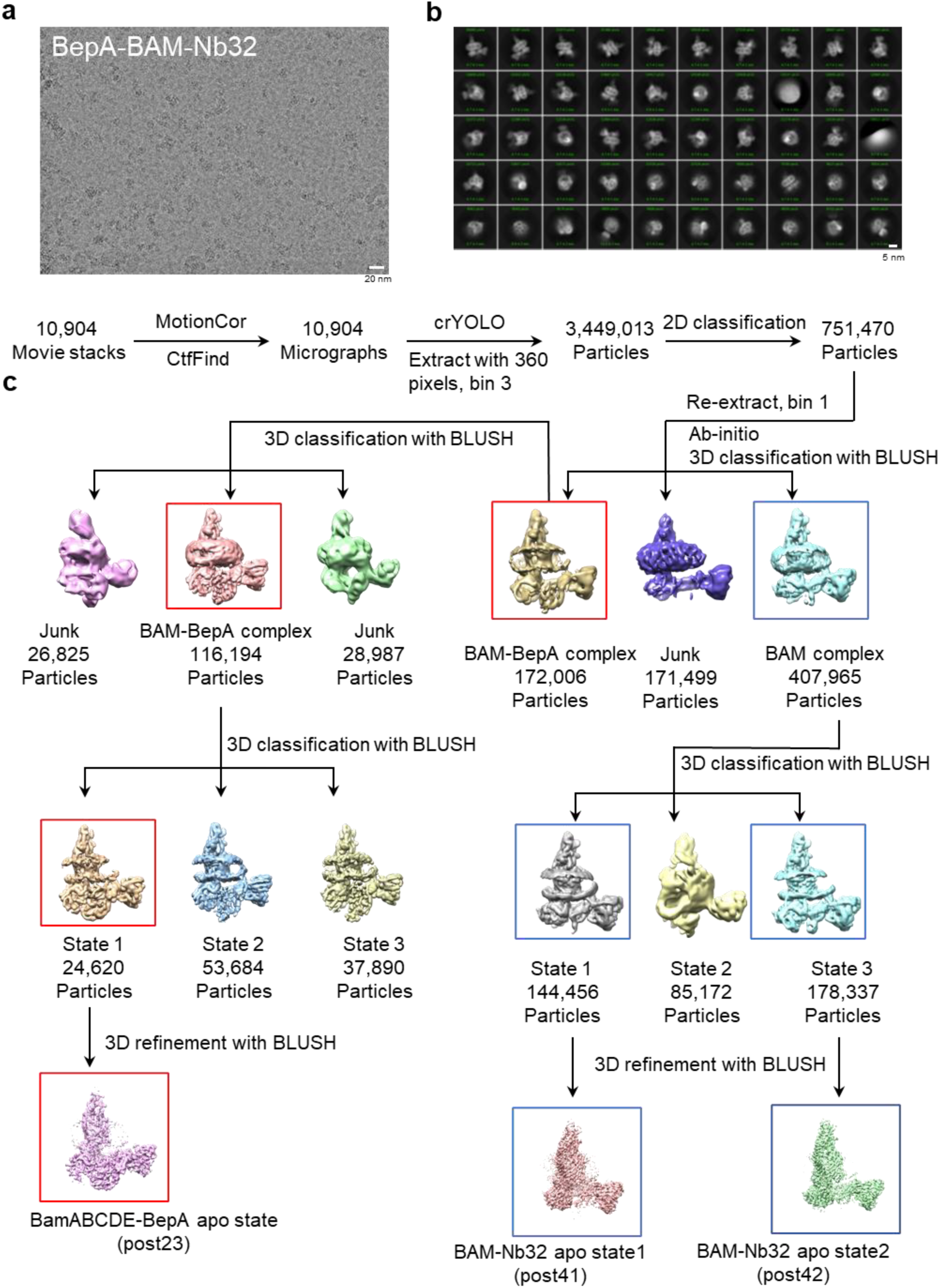
Purification and cryoEM 3D reconstruction of BepA-BAM-Nb32 complex. (**a**) Raw cryoEM micrograph BepA-BAM-Nb32 collected at 300kV and 60k magnification. (**b**) 2D classes obtained for BepA-BAM-Nb32 complex. (**c**) Image processing and structure determination pipeline used for the BepA-BAM-Nb32 dataset.

**Extended Data Figure 6.**
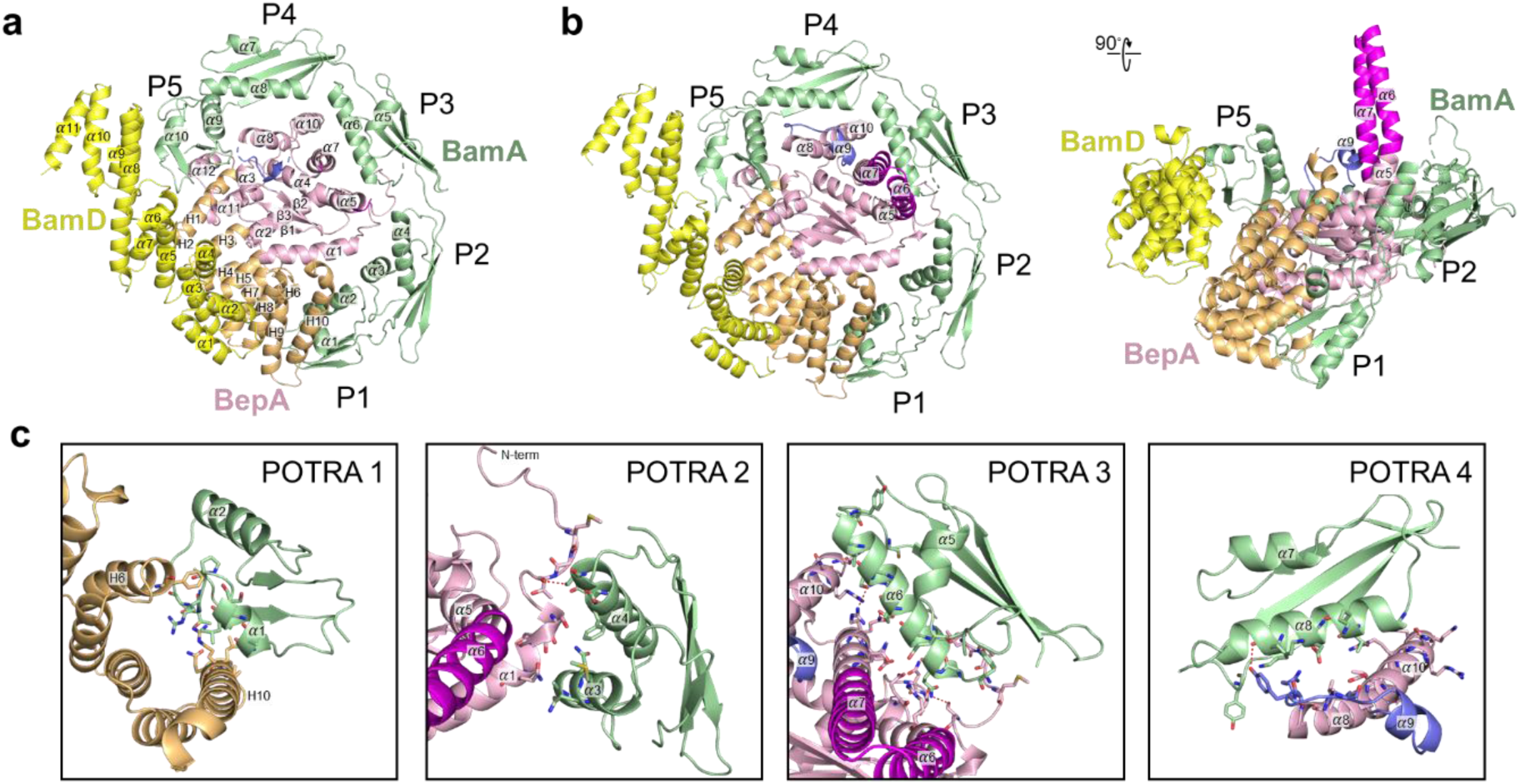
Structural analysis of the BepA-BamA contact. (**a, b**) Top view of the BepA-BAM-Nb32 complex (a) and top and side view of the BepA-BAM-Nb5 complex (b). Figures concentrate on contact regions, showing BamD (yellow), BamA POTRA 1-5 (green) and BepA. The BepA M48 metalloprotease domain is colored pink (α1-α12) and its TPR domain is colored orange (H1-H10). The AI-plug and α6-lid are colored slate blue and magenta, respectively. BepA adopts identical contacts in both complexes, with the BepA TPR in a 140.3 Å^2^ contact with BamD, and 463.8 Å^2^ contact with BamA P1, and its M48 metalloprotease domain in a 413.7 Å^2^ contact with BamA P2, 986.0 Å^2^ contact with BamA P3, and 385.1 Å^2^ contact with BamA P4. (**c**) Close-up views of the BepA interaction with BamA POTRA 1-4 as found in the BepA-BAM-Nb5 complex. Nb5 binds the N-terminal tip of POTRA 1 (not shown).

**Extended Data Figure 7.**
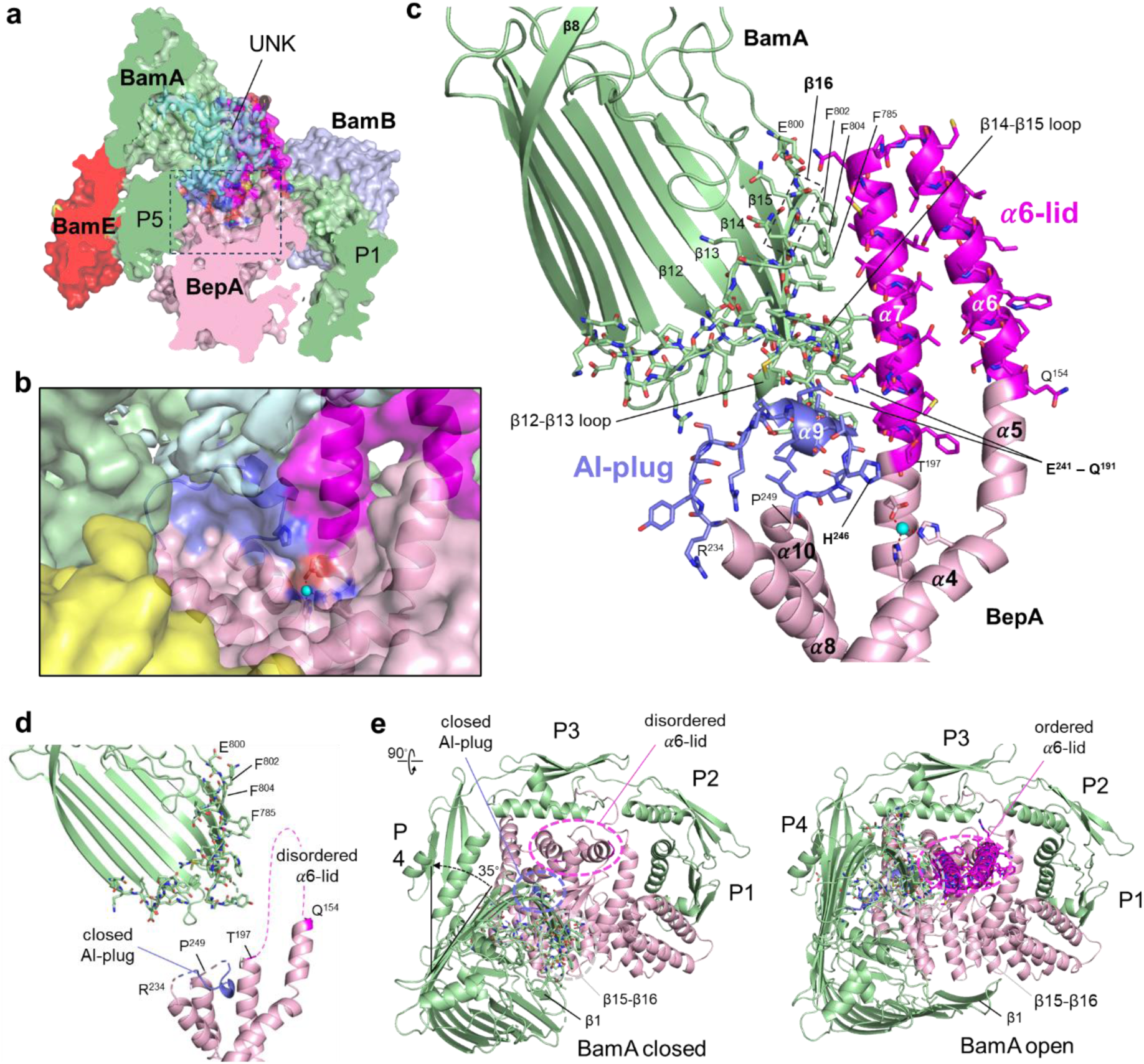
Structural analysis of the BepA-BAM complexes. (**a, b**) Slice-through and zoomed view (b, boxed area, showing catalytic site) of the BepA-BAM-Nb5 complex (*i.e.* BamABCDE-BepA-OMP-UNK) shown in molecular surface representation, superimposed with the cryoEM density of the unidentified substrate protein (light cyan, labeled UNK) in the lumen of the BamA β-barrel. BamA, BamB, BamD and BamE are colored green, light blue, yellow and red, respectively. BepA is colored pink, with the AI-plug and α6-lid colored slate blue and magenta, respectively. Zn^2+^ (cyan sphere) ligands are shown in stick representation. (**c**) Cartoon representation of the BepA-BAM-Nb5 complex, shown in side view and focused on the interaction zone of the BepA catalytic site (α4-α10), AI-plug (slate blue, residues R^234^ to P^249^), α6-lid (magenta, residues Q^154^ to T^197^) and the BamA β-barrel (shown from β8-β16). BepA M48 metalloprotease domain is in a 673.4 Å^2^ contact area with the β-barrel of BamA. (**d**) Cartoon representation of the BepA-BAM-Nb32 complex, shown in equivalent view as BepA-BAM-Nb5 in panel c. The α6-lid and α6-α7 loop (*i.e.* residues Q^154^ to T^197^) are not resolved in the cryoEM density, the AI-plug (α9; residues R^234^ to P^249^) and loops connecting it to α8 and α10 are only partially resolved in the cryoEM density. (**e**) Top views of the BepA-BAM-Nb32 (left) and BepA-BAM-Nb5 (right) complexes, showing BamA (green) and BepA (pink); other BAM subunits are not shown for clarity. Shown in stick are residues forming the contact zones of the BepA α6-lid and AI-plug with the BamA β-barrel and periplasmic loops (as in panels c, d). Opening of the BamA β-barrel during OMP insertion (OMP not shown) is accompanied with a ∼35° rotation relative to the POTRA domains, and exposes a contact surface in BamA β15-16 that binds and orders the membrane-inserted α6-lid of BepA.

**Extended Data Figure 8.**
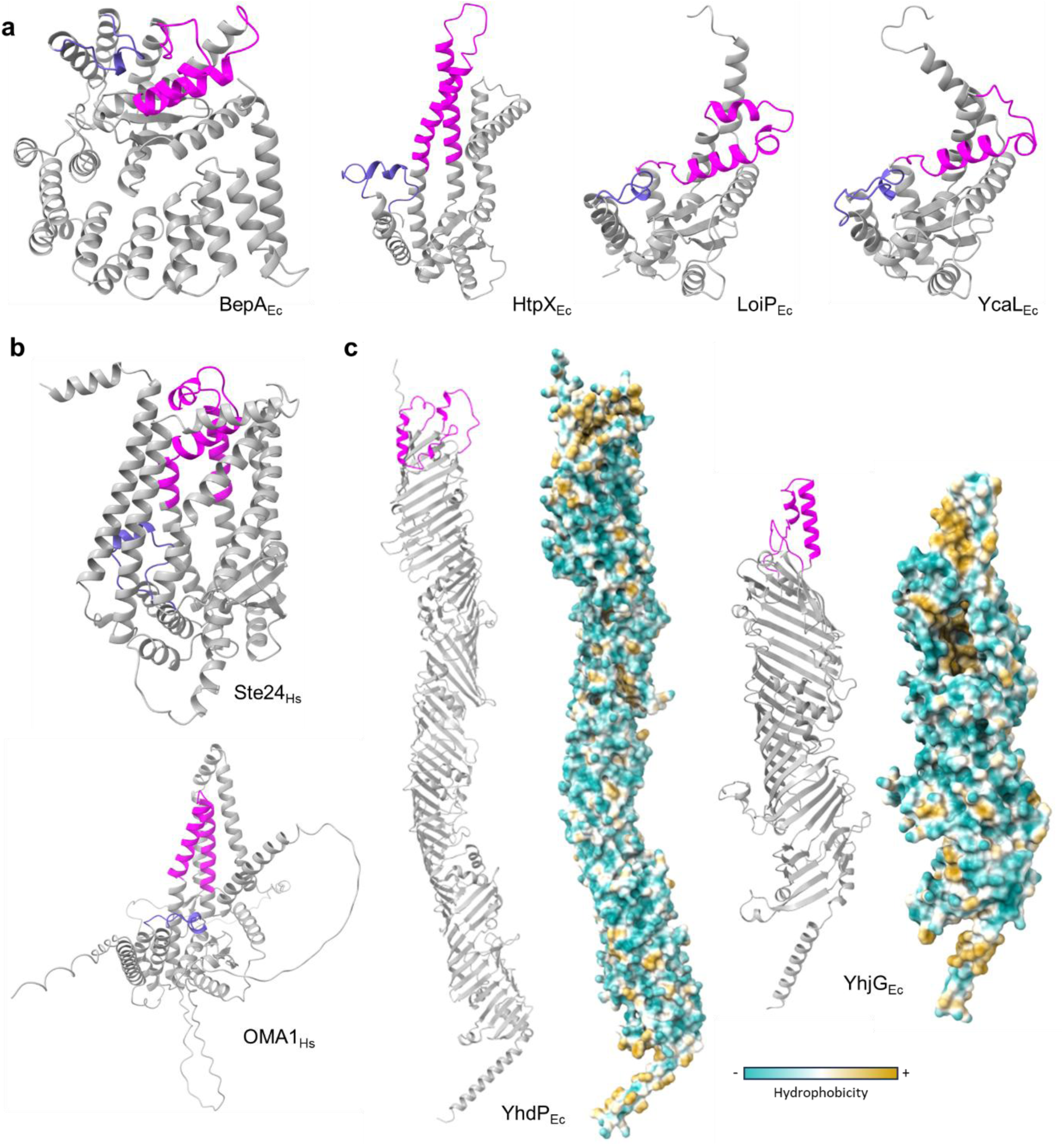
Structural comparisons of M48 metalloproteases and AsmA-like proteins. AlphaFold3 structure predictions are shown in cartoon representation. The AI-plug and α6-lid of BepA in a close-conformation are colored slate blue and magenta, respectively. These conserved structural features found in M48 metalloprotease homologues of (**a**) *Escherichia coli* and (**b**) *Homo sapiens* are highlighted in the same colors. (**c**) The hydrophobic α6-lid-like structures of the AsmA-like proteins YhdP and YhjG of *Escherichia coli* are colored magenta. The surface representations of YhdP and YhjG structures are colored by hydrophobicity (from blue for the most hydrophilic, to white, to yellow for the most hydrophobic). Ec, *Escherichia coli*; Hs, *Homo sapiens*.

**Supplementary Table S1.**
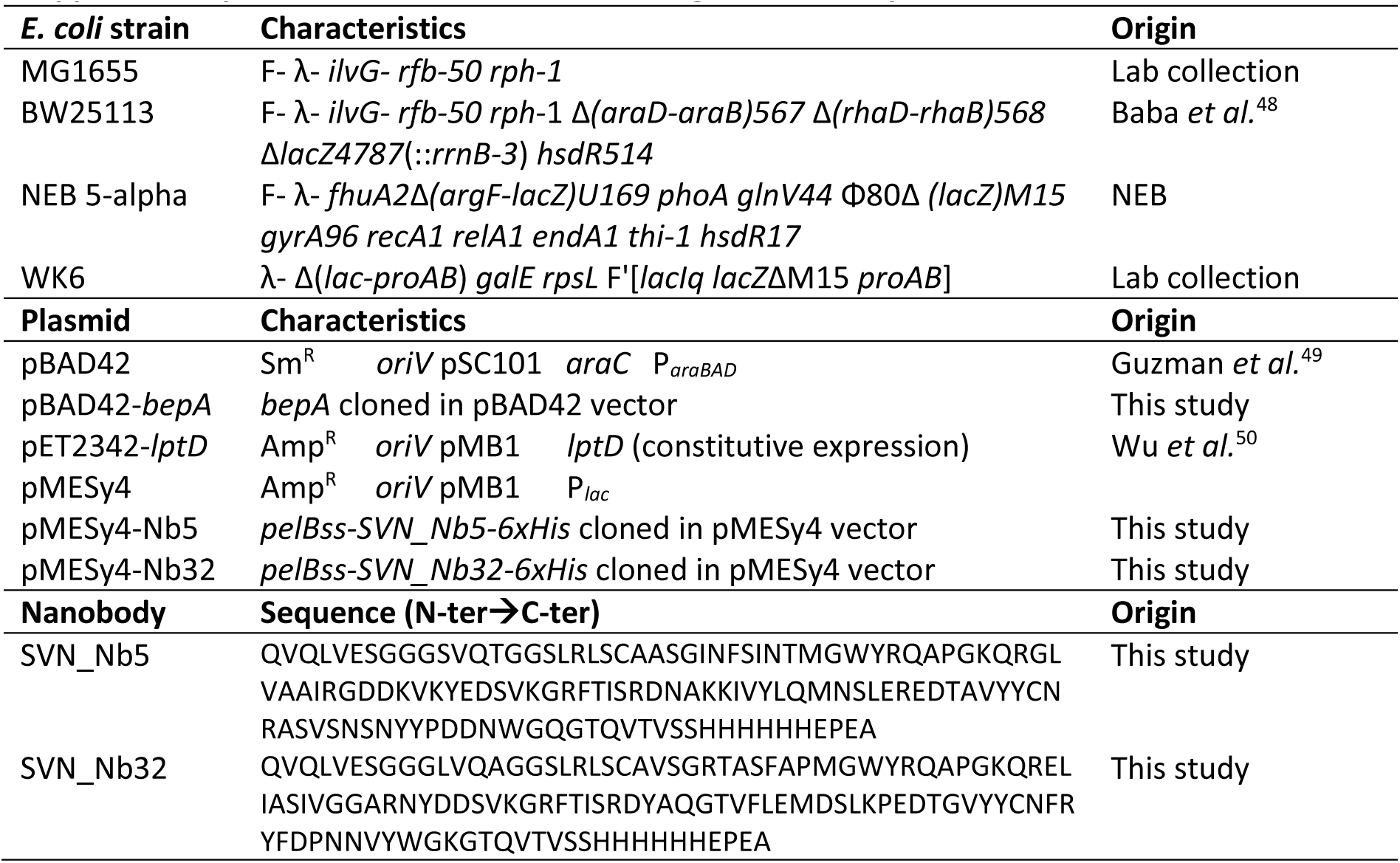
Characteristics and origin of strains, plasmids, and nanobodies.

**Supplementary Table S2.** Cryo-EM data collection, refinement, and validation statistics.

|  | BepA-BAM-nb32<br>EMD-55064<br>PDB: 9SOX | BepA-BAM-nb5<br>EMD-55063<br>PDB: 9SOW |
| --- | --- | --- |
| **Data collection and processing** |  |  |
| Magnification | 60 000 | 60 000 |
| Voltage (kV) | 300 | 300 |
| Electron exposure (e<sup>-</sup>/Å<sup>2</sup>) | 60 | 60 |
| Defocus range (μm) | 0.8-2.0 | 0.8-2.0 |
| Pixel size (Å) | 0.76 | 0.76 |
| Symmetry imposed | C1 | C1 |
| Initial particle images (no.) | 3 449 013 | 10 108 143 |
| Final particle images (no.) | 24 620 | 174 705 |
| Map resolution (Å) | 6.36 | 3.23 |
| FSC threshold | 0.143 | 0.143 |
| Map resolution range (Å) | 5.1-7.8 | 2.1-3.7 |
| **Model** |  |  |
| Initial model used (PDB code) | 9CNW; 6AIT | 7YE4; 6AIT |
| Model resolution (Å) | 6.4 | 3.23 |
| FSC threshold | 0.143 | 0.143 |
| Model composition |  |  |
| Chains | 7 | 8 |
| Non-hydrogen atoms | 16384 | 15079 |
| Protein residues | 2099 | 1943 |
| Ligands (non-standard residues) | 1 | 1 |
| R.m.s. deviations |  |  |
| Bond lengths (Å) | 0.004 | 0.003 |
| Bond angles (°) | 0.738 | 0.823 |
| Validation |  |  |
| MolProbity score | 2.17 | 2.12 |
| Clashscore | 16.93 | 12.14 |
| Poor rotamers (%) | 0 | 1.51 |
| Ramachandran plot |  |  |
| Favored (%) | 93.25 | 94.24 |
| Allowed (%) | 6.25 | 5.7 |
| Disallowed (%) | 0 | 0.05 |
| **Model vs Data** |  |  |
| CC (mask) | 0.76 | 0.68 |
| CC (box) | 0.88 | 0.79 |
| CC (peaks) | 0.57 | 0.68 |
| CC (volume) | 0.73 | 0.63 |

## REFERENCES

1. Silhavy, T. J., Kahne, D. & Walker, S. The bacterial cell envelope. Cold Spring Harb. Perspect. Biol. 2, a000414 (2010).

2. Zgurskaya, H. I., López, C. A. & Gnanakaran, S. Permeability Barrier of Gram-Negative Cell Envelopes and Approaches To Bypass It. ACS Infect. Dis. 1, 512–522 (2015).

3. Rojas, E. R. et al. The outer membrane is an essential load-bearing element in Gram-negative bacteria. Nat. 2018 5597715 559, 617–621 (2018).

4. Deghelt, M. et al. Peptidoglycan-outer membrane attachment generates periplasmic pressure to prevent lysis in Gram-negative bacteria. Nat. Microbiol. 10, 1963–1974 (2025).

5. Combs, A. N. & Silhavy, T. J. Periplasmic Chaperones: Outer Membrane Biogenesis and Envelope Stress. Annu. Rev. Microbiol. 78, 191–211 (2024).

6. Doyle, M. T. & Bernstein, H. D. Function of the Omp85 Superfamily of Outer Membrane Protein Assembly Factors and Polypeptide Transporters. Annu. Rev. Microbiol. 76, 259–279 (2022).

7. Tomasek, D. et al. Structure of a nascent membrane protein as it folds on the BAM complex. Nature 583, 473–478 (2020).

8. Wang, Q. et al. Structural insight into the SAM-mediated assembly of the mitochondrial TOM core complex. Science 373, 1377–1381 (2021).

9. Tomasek, D. & Kahne, D. The assembly of β-barrel outer membrane proteins. Curr. Opin. Microbiol. 60, 16–23 (2021).

10. Doyle, M. T. et al. Cryo-EM structures reveal multiple stages of bacterial outer membrane protein folding. Cell 185, 1143–1156.e13 (2022).

11. Shen, C. et al. Structural basis of BAM-mediated outer membrane β-barrel protein assembly. Nature 617, 185–193 (2023).

12. Cho, S. H. et al. Detecting Envelope Stress by Monitoring β-Barrel Assembly. Cell 159, 1652–1664 (2014).

13. Soltes, G. R., Martin, N. R., Park, E., Sutterlin, H. A. & Silhavy, T. J. Distinctive Roles for Periplasmic Proteases in the Maintenance of Essential Outer Membrane Protein Assembly. J. Bacteriol. 199, (2017).

14. Combs, A. N. & Silhavy, T. J. The sacrificial adaptor protein Skp functions to remove stalled substrates from the β-barrel assembly machine. Proc. Natl. Acad. Sci. U. S. A. 119, (2022).

15. Dartigalongue, C., Missiakas, D. & Raina, S. Characterization of the Escherichia coli σE Regulon. J. Biol. Chem. 276, 20866–20875 (2001).

16. Walsh, N. P., Alba, B. M., Bose, B., Gross, C. A. & Sauer, R. T. OMP peptide signals initiate the envelope-stress response by activating DegS protease via relief of inhibition mediated by its PDZ domain. Cell 113, 61–71 (2003).

17. Rhodius, V. A., Suh, W. C., Nonaka, G., West, J. & Gross, C. A. Conserved and variable functions of the sigmaE stress response in related genomes. PLoS Biol. 4, 0043–0059 (2006).

18. Narita, S. I., Masui, C., Suzuki, T., Dohmae, N. & Akiyama, Y. Protease homolog BepA (YfgC) promotes assembly and degradation of β-barrel membrane proteins in Escherichia coli. Proc. Natl. Acad. Sci. U. S. A. 110, 3612–3621 (2013).

19. Daimon, Y. et al. The TPR domain of BepA is required for productive interaction with substrate proteins and the β-barrel assembly machinery complex. Mol. Microbiol. 106, 760–776 (2017).

20. Shahrizal, M. et al. Structural Basis for the Function of the β-Barrel Assembly-Enhancing Protease BepA. J. Mol. Biol. 431, 625–635 (2019).

21. Bryant, J. A. et al. Structure-Function Characterization of the Conserved Regulatory Mechanism of the Escherichia coli M48 Metalloprotease BepA. J. Bacteriol. 203, e00434–20 (2020).

22. Akiyama, Y. Quality control of cytoplasmic membrane proteins in Escherichia coli. J. Biochem. 146, 449–454 (2009).

23. Käser, M., Kambacheld, M., Kisters-Woike, B. & Langer, T. Oma1, a Novel Membrane-bound Metallopeptidase in Mitochondria with Activities Overlapping with the m-AAA Protease. J. Biol. Chem. 278, 46414–46423 (2003).

24. Head, B., Griparic, L., Amiri, M., Gandre-Babbe, S. & Van Der Bliek, A. M. Inducible proteolytic inactivation of OPA1 mediated by the OMA1 protease in mammalian cells. J. Cell Biol. 187, 959–966 (2009).

25. Pryor, E. E. et al. Structure of the integral membrane protein CAAX protease Ste24p. Science 339, 1600–1604 (2013).

26. Quigley, A. et al. The structural basis of ZMPSTE24-dependent laminopathies. Science 339, 1604–1607 (2013).

27. Baker, M. J. et al. Stress-induced OMA1 activation and autocatalytic turnover regulate OPA1-dependent mitochondrial dynamics. EMBO J. 33, 578 (2014).

28. Ast, T., Michaelis, S. & Schuldiner, M. The protease Ste24 clears clogged translocons. Cell 164, 103 (2016).

29. Daimon, Y. et al. Reversible autoinhibitory regulation of Escherichia coli metallopeptidase BepA for selective β-barrel protein degradation. Proc. Natl. Acad. Sci. U. S. A. 117, 27989–27996 (2020).

30. Miyazaki, R., Watanabe, T., Yoshitani, K. & Akiyama, Y. Edge-strand of BepA interacts with immature LptD on the β-barrel assembly machine to direct it to on-and off-pathways. Elife 10, (2021).

31. Paradis-Bleau, C., Kritikos, G., Orlova, K., Typas, A. & Bernhardt, T. G. A Genome-Wide Screen for Bacterial Envelope Biogenesis Mutants Identifies a Novel Factor Involved in Cell Wall Precursor Metabolism. PLOS Genet. 10, e1004056 (2014).

32. Lazar, S. W. & Kolter, R. SurA assists the folding of Escherichia coli outer membrane proteins. J. Bacteriol. 178, 1770–1773 (1996).

33. Rouvière, P. E. & Gross, C. A. SurA, a periplasmic protein with peptidyl-prolyl isomerase activity, participates in the assembly of outer membrane porins. Genes Dev. 10, 3170–3182 (1996).

34. Ades, S. E., Connolly, L. E., Alba, B. M. & Gross, C. A. The Escherichia coli σE-dependent extracytoplasmic stress response is controlled by the regulated proteolysis of an anti-σ factor. Genes Dev. 13, 2449 (1999).

35. Fenn, K. L. et al. Outer membrane protein assembly mediated by BAM-SurA complexes. Nat. Commun. 15, 1–17 (2024).

36. Sochacki, K. A., Shkel, I. A., Record, M. T. & Weisshaar, J. C. Protein Diffusion in the Periplasm of E. coli under Osmotic Stress. Biophys. J. 100, 22 (2011).

37. Pilizota, T. & Shaevitz, J. W. Plasmolysis and cell shape depend on solute outer-membrane permeability during hyperosmotic shock in E. coli. Biophys. J. 104, 2733–2742 (2013).

38. Kumar, S. & Ruiz, N. Bacterial AsmA-Like Proteins: Bridging the Gap in Intermembrane Phospholipid Transport. *Contact (Thousand Oaks (Ventura County*, Calif.)) 6, (2023).

39. Cooper, B. F. et al. Phospholipid Transport Across the Bacterial Periplasm Through the Envelope-spanning Bridge YhdP. J. Mol. Biol. 437, 168891 (2025).

40. Hackbarth, C. J. & Chambers, H. F. blaI and blaR1 regulate beta-lactamase and PBP 2a production in methicillin-resistant Staphylococcus aureus. Antimicrob. Agents Chemother. 37, 1144–1149 (1993).

41. Alexander, J. A. N. et al. Structural basis of broad-spectrum β-lactam resistance in Staphylococcus aureus. Nature 613, 375–382 (2023).

42. Wai, T. et al. Imbalanced OPA1 processing and mitochondrial fragmentation cause heart failure in mice. Science 350, (2015).

43. Rivera-Mejías, P. et al. The mitochondrial protease OMA1 acts as a metabolic safeguard upon nuclear DNA damage. Cell Rep. 42, (2023).

44. Chen, L. et al. Inhibition of mitochondrial OMA1 ameliorates osteosarcoma tumorigenesis. Cell Death Dis. 15, (2024).

45. Barrowman, J., Wiley, P. A., Hudon-Miller, S. E., Hrycyna, C. A. & Michaelis, S. Human ZMPSTE24 disease mutations: residual proteolytic activity correlates with disease severity. Hum. Mol. Genet. 21, 4084–4093 (2012).

46. Fu, B., Wang, L., Li, S. & Dorf, M. E. ZMPSTE24 defends against influenza and other pathogenic viruses. J. Exp. Med. 214, 919–929 (2017).

47. Shilagardi, K., Spear, E. D., Abraham, R., Griffin, D. E. & Michaelis, S. The Integral Membrane Protein ZMPSTE24 Protects Cells from SARS-CoV-2 Spike-Mediated Pseudovirus Infection and Syncytia Formation. MBio 13, e02543–22 (2022).

48. Baba, T. et al. Construction of *Escherichia coli* K-12 in-frame, single-gene knockout mutants: the Keio collection. Mol. Syst. Biol. 2, (2006).

49. Guzman, L. M., Belin, D., Carson, M. J. & Beckwith, J. Tight regulation, modulation, and high-level expression by vectors containing the arabinose PBAD promoter. J. Bacteriol. 177, 4121–4130 (1995).

50. Wu, T. et al. Identification of a protein complex that assembles lipopolysaccharide in the outer membrane of Escherichia coli. Proc. Natl. Acad. Sci. U. S. A. 103, 11754–11759 (2006).

51. Datsenko, K. A. & Wanner, B. L. One-step inactivation of chromosomal genes in Escherichia coli K-12 using PCR products. Proc. Natl. Acad. Sci. U. S. A. 97, 6640–6645 (2000).

52. Janssens, A. et al. SlyB encapsulates outer membrane proteins in stress-induced lipid nanodomains. Nature 626, 617–625 (2024).

53. Pardon, E. et al. A general protocol for the generation of Nanobodies for structural biology. Nat. Protoc. 9, 674–693 (2014).

54. Mastronarde, D. N. Automated electron microscope tomography using robust prediction of specimen movements. J. Struct. Biol. 152, 36–51 (2005).

55. Zheng, S. Q. et al. MotionCor2: Anisotropic correction of beam-induced motion for improved cryo-electron microscopy. Nat. Methods 14, 331–332 (2017).

56. Rohou, A. & Grigorieff, N. CTFFIND4: Fast and accurate defocus estimation from electron micrographs. J. Struct. Biol. 192, 216–221 (2015).

57. Wagner, T. et al. SPHIRE-crYOLO is a fast and accurate fully automated particle picker for cryo-EM. *Commun*. Biol. 2, 1–13 (2019).

58. Zivanov, J. et al. New tools for automated high-resolution cryo-EM structure determination in RELION-3. Elife 7, (2018).

59. Punjani, A., Rubinstein, J. L., Fleet, D. J. & Brubaker, M. A. CryoSPARC: Algorithms for rapid unsupervised cryo-EM structure determination. Nat. Methods 14, 290–296 (2017).

60. Pettersen, E. F. et al. UCSF Chimera--a visualization system for exploratory research and analysis. J. Comput. Chem. 25, 1605–1612 (2004).

61. Emsley, P. & Cowtan, K. Coot: model-building tools for molecular graphics. Acta Crystallogr. D. Biol. Crystallogr. 60, 2126–2132 (2004).

62. Afonine, P. V. et al. Real-space refinement in PHENIX for cryo-EM and crystallography. *Acta Crystallogr. Sect. D, Struct*. Biol. 74, 531–544 (2018).

63. Pettersen, E. F. et al. UCSF ChimeraX: Structure visualization for researchers, educators, and developers. Protein Sci. 30, 70–82 (2021).

